# Allele-resolved monosome and polysome sequencing identifies functional cis-acting variants affecting mRNA translation efficiency

**DOI:** 10.64898/2026.05.07.723424

**Authors:** Laura Alunno, Ilaria Massignani, Meriem Hadjer Hamadou, Fabio Mazza, Daniele Peroni, Romina Belli, Erik Dassi, Alessandro Romanel, Alberto Inga

## Abstract

To prioritize germline genetic variants affecting mRNA fate at the post-transcriptional and translational levels, we leveraged sucrose-gradient-based isolation of 80S monosomes and polysomes, followed by mRNA retrieval and paired-end sequencing. Total cytoplasmic RNA was also sequenced for comparison. Experiments were performed in the non-transformed cell line RPE-1, cultured under basal conditions or upon p53 activation by Nutlin. Differential gene expression analysis confirmed a canonical p53 response. Heterozygous SNPs and SNVs were identified from the RNA-seq data, and allelic fractions (AF) were calculated for total, monosomal, and polysomal mRNAs. Variants showing reproducible AF differences across fractions beyond experimental variability were defined as tranSNPs. Among nearly 7000 heterozygous variants analyzable in polysomal or total RNA and over 5000 in monosomal mRNA, 1155 displayed a significant imbalance. Reporter assays performed in both RPE-1 and HCT116 cells validated allelic or haplotype effects for 17 selected variants in UTRs and coding regions, confirming differences in 15 cases, with evidence of cell line-specific responses. Proteomic analysis further supported allelic imbalance for selected missense variants. Overall, tranSNPs were identified in a non-transformed cell line at frequencies comparable to those in cancer cells, thereby extending their implications in human physiology. Further, monosome profiling enabled improved detection sensitivity of tranSNPs without positional bias, suggesting that 80S profiling improves detection of allele-specific translational regulation in RPE-1 cells.

**Graphical Abstract:** 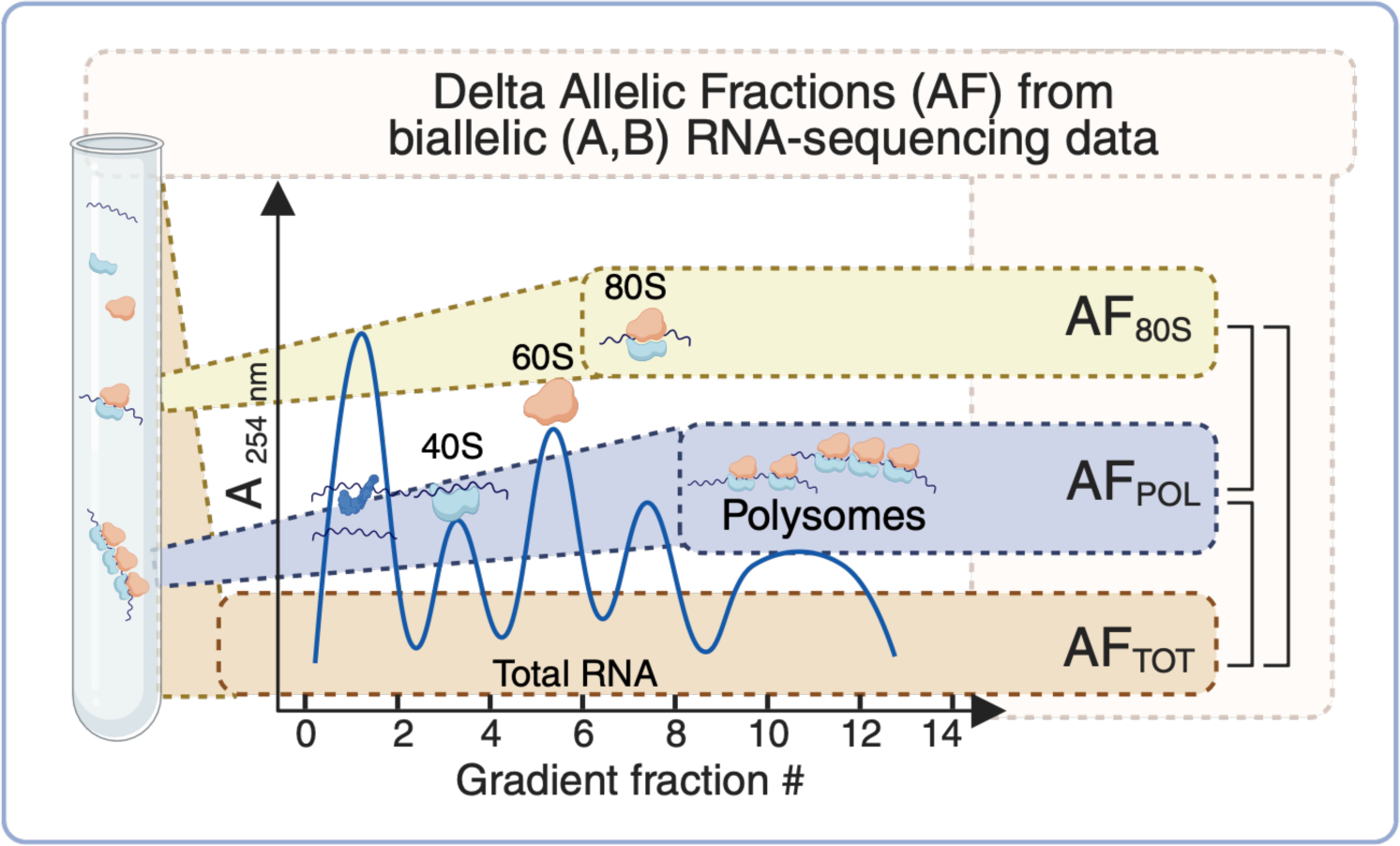

## Introduction

The relevance of the post-transcriptional and translational controls of gene expression has recently been emphasized (Mata et al. 2005; Mitschka and Mayr 2022; Wu and Bazzini 2023; Biffo et al. 2024; Orellana et al. 2022; Zhao et al. 2016; Robichaud et al. 2019; Ruggero 2013; Jia et al. 2020). However, only a limited number of studies have explored genetic variants affecting post-transcriptional events (Robert and Pelletier 2018; Huang et al. 2025; Li and Chen 2023; Zheng et al. 2025). Furthermore, an established experimental pipeline for systematically prioritize functional variants that affect mRNA fate is lacking, although tools using, for example, global translation efficiency data and machine learning to reach nucleotide resolution are emerging (Zheng et al. 2025; Xiong et al. 2025).

We recently developed a method for the identification of tranSNPs, where tranSNPs were defined as “a class of functional SNPs affecting the mRNA translation potential” (Valentini et al. 2021). The approach relies on the comparative analysis of the allelic fraction (AF) of the alternative allele across polysome-bound mRNAs (POL) and total mRNAs (TOT) for the SNPs that are heterozygous and transcribed in a given cell model. Polysome-bound mRNAs are considered a proxy for the actively translated mRNAs, while total mRNAs represent a proxy for less translated mRNAs. When two SNP alleles are significantly imbalanced in polysomal RNA compared to total RNA, one allele is evidently enriched in the polysomes, hence the SNP is identified as a putative tranSNP. The pipeline was applied to the breast cancer cell line MCF7, and 147 SNPs, corresponding to about 4% of the analyzable SNPs, were identified as putative tranSNPs. No enrichment for specific pathways was observed among the genes harboring the tranSNPs. About 25% of tranSNPs were located in the UTRs, with the remainder in coding regions.

The pipeline was then applied to a dataset based on total and polysomal RNA sequencing of the colon cancer cell line HCT116, consisting of four biological replicates, but with lower coverage than the MCF7 dataset (Hamadou et al. 2025). A total of 40 putative tranSNPs, corresponding to approximately 7% of the analyzable SNPs, were identified. As in the MCF7 dataset, no enrichment for specific pathways or gene networks was observed among the putative tranSNPs. After a first step of validation through Sanger sequencing on new biological replicates and luciferase reporter assays, two tranSNPs were selected for an in-depth validation: rs1053639 on DDIT4 3’ UTR and the rs1554710467 missense variant on EIF4H coding sequence (CDS), the latter one validated also by allele-specific proteomics (Hamadou et al. 2026).

In this study, we further implemented the tranSNP workflow and applied it to the human retinal pigment epithelial RPE-1 cell line to explore the tranSNP phenomenon in non-cancer cells. RPE-1 cells were selected as representatives of non-cancer cell lines because of their established use in research, transfection, and editing amenability, as well as their well-described and stable karyotype (Volpe et al. 2025). The RPE-1 dataset was generated using high coverage (on average ∼130 million reads) paired-end sequencing of three biological replicates. The high coverage enabled the identification of a large number of putative tranSNPs, even within the same mRNA, and allowed a broader exploration of the tranSNP effect, exploring haplotype structure and mechanisms. Importantly, an additional fraction (80S), corresponding to the mRNAs bound to the 80S monosome, was analyzed and compared to the well-established polysome-bound and total cytoplasmic mRNA fractions in the mock condition. Since the translation status of the 80S monosome is still debated (Blandy et al. 2025; Heyer and Moore 2016), the comparison of polysomal and total mRNAs with the 80S-bound mRNAs holds the potential to contribute to the field by clarifying the potential of the 80S, and to diminish the potential bias originated by the incorporation of polysome-bound mRNAs in total mRNAs, by focusing on a fraction that is free from polysomes.

## Results

### TranSNPs can be identified in non-cancer cells

To follow up on the potential relevance of tranSNPs, we wondered whether the allelic imbalance leading to the identification of tranSNPs was exclusive or particularly prominent in cancer cells due to their broadly rewired cellular deregulation, or whether it also occurred in non-transformed cells. We generated a new catalog of tranSNPs in the hTERT-immortalized retinal pigment epithelial cells (RPE-1) characterized by high coverage (average 134 million reads, **Table S1**) and three biological replicates for each fraction. In addition to the mock condition, the treatment with 10 µM Nutlin for 16 hours was applied. Differential gene expression analyses confirmed that the treatment activated the p53 and cell cycle arrest pathways (**Figure S1**). Nutlin treatment through MDM2 inhibition and p53 activation is known to elicit broad changes at transcriptional but also post-transcriptional and translational regulation levels (Tiu et al. 2021; Rizzotto et al. 2020).

We identified heterozygous SNPs from RNA sequencing data by applying a minimum coverage threshold of 20 reads per replicate and condition. Additionally, we filtered variants based on allelic fractions ranging from 20% to 80% (**Table S1**). Then, we calculated the allelic imbalance across polysome-bound and total cytoplasmic mRNAs (delta allelic fraction, dAF) for analyzable heterozygous SNPs, and used nominal p-values derived from paired t-tests across biological replicates to prioritize and catalog putative RPE-1 tranSNPs. For these dAF calculations, cases in which AF was within the 20%-80% range in only one fraction were included, to avoid missing extreme imbalances. To improve the tranSNP identification pipeline, we added a further fraction to this dataset, represented by 80S monosome-bound mRNAs, and compared it with total and polysome-bound mRNAs. We identified 411 tranSNPs in RPE-1 by comparing polysome-bound and total mRNAs (POL-TOT), 359 by comparing polysome-bound and 80S-bound mRNAs (POL-80S), and 299 by comparing the 80S-bound and total mRNAs (80S-TOT), indicating that tranSNPs can be identified in non-cancer settings as well (**Table S1**; **Figure S2**). We also applied an additional filter of −2SD > dAF > 2SD to the identified tranSNPs, based on the variance in AF measurements across all analyzable heterozygous SNPs for each condition or treatment, to reduce false positives and prioritize variants with larger allelic imbalance. This approach complements nominal p-value-based prioritization by incorporating orthogonal features, including effect size and consistency across replicates, thereby increasing stringency. With this approach, we restricted the tranSNP lists to 73 when comparing polysome-bound and total mRNAs (POL-TOT), 107 when comparing polysome-bound and 80S-bound mRNAs (POL-80S), and 56 when comparing the 80S-bound and total mRNAs (80S-TOT). In total, 1155 unique tranSNPs were found in 5’UTR (78), 3’UTR (462), and coding regions (615).

### The use of 80S-bound mRNAs improves resolution in the identification of tranSNPs

The translational status of the 80S monosome is still highly debated in the field (Blandy et al. 2025; Heyer and Moore 2016). We exploited the RPE-1 dataset to evaluate the allelic imbalance (dAF, delta allelic fraction) of tranSNPs, focusing on the 80S-monosome data for the comparisons. By comparing the correlation between allelic fractions in the 80S-TOT and POL-80S datasets, expressed as R^2^ scores, we found that the POL-80S comparison better discriminated differences in allelic fraction imbalances (**Figure 1A-D, Figure S2**). Notably, most of the tranSNPs from 80S-bound mRNAs displayed complementary delta allelic imbalance in the 80S-TOT or POL-80S comparisons **(Figure 1E)**. This systematic observation led us to think that the 80S fraction and the polysomal fraction represent extreme conditions for dAF measurements, while TOT-POL dAF values (even when statistical significance was not achieved) laid in the middle. Interestingly, we found that the percentage of tranSNPs relative to the analyzable SNPs was higher in the 80S-POL (about 6.5% based on nominal p-values, or 1.9% after applying variance-based filtering) than in 80S-TOT (about 5.4% or 1%) or POL-TOT (5.5% or 1%) comparisons. This data further supported the hypothesis that 80S monosomes are a better proxy for unmasking allelic differences in mRNA translation compared to total cytoplasmic mRNAs, at least in RPE-1 cells, providing higher resolution in the identification of tranSNPs through our pipeline. Although limited by potential heterogeneities in the overall composition and complexity of the fractions, we explored differential gene expression changes among fractions (**Figure S3**). The strongest transcriptional separation was observed when comparing the 80S/monosomal and polysome-associated fractions, whereas the other comparisons showed less pronounced differences. Functional enrichment analysis of fraction-associated DEGs revealed a prominent enrichment for cytoplasmic translation and structural constituents of the ribosome in the 80S fraction compared with polysomes. Similarly, ribosomal protein transcripts were upregulated in the comparison between total and polysomal mRNAs both in the mock and nutlin-treated conditions. These data are consistent with the observation that TOP mRNAs have lower translation efficiency or can be translated by monosomes more than polysomes (Jia et al. 2021; Thoreen et al. 2012; Schneider et al. 2022).

**Figure 1.**
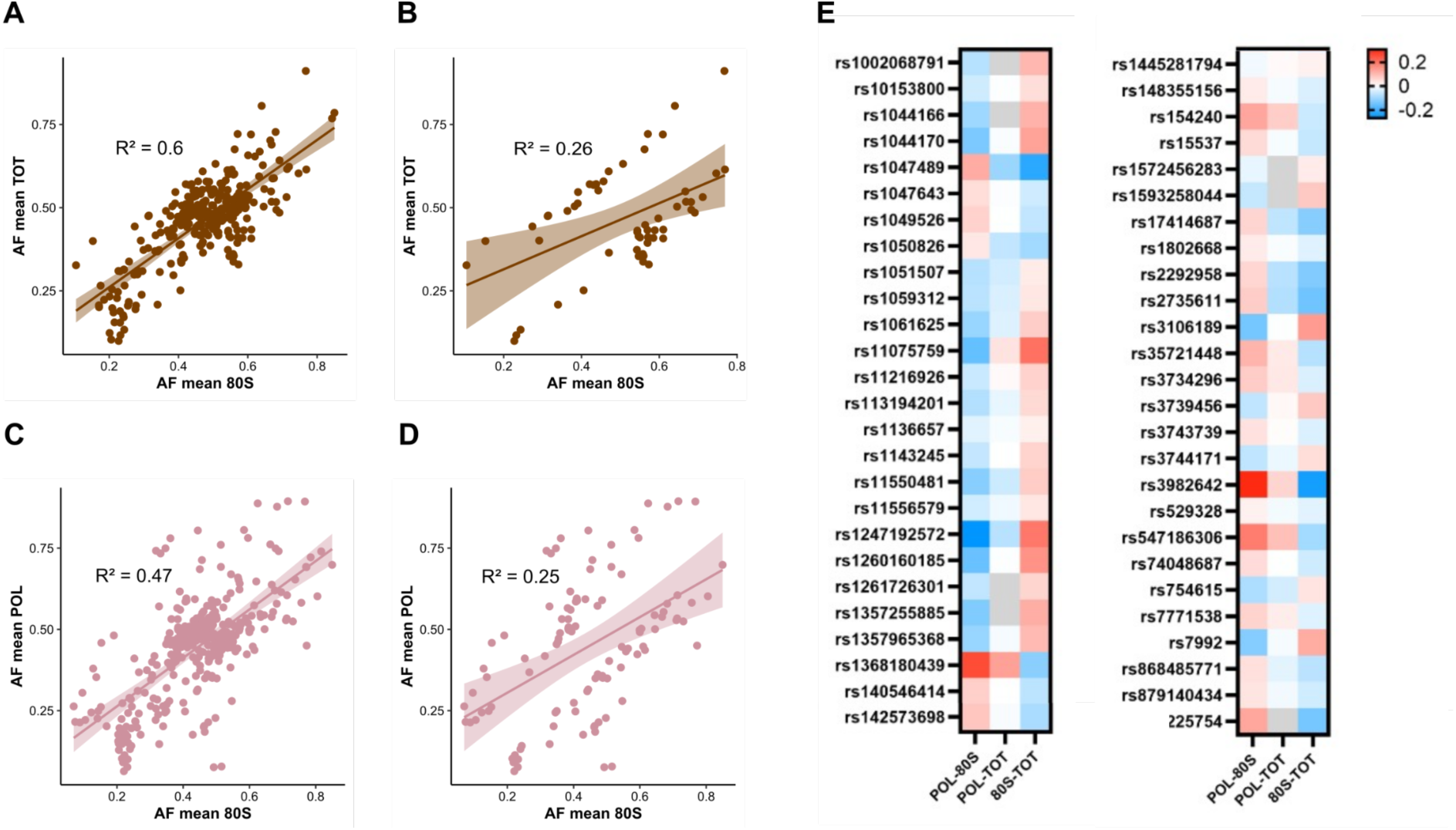
The comparison of polysome-bound and 80S-bound mRNAs improves discrimination in tranSNPs discrimination. (A) Distribution of allelic fraction (AF) values across 80S and TOT mRNA fractions for the tranSNPs selected based on p < 0.05. (B) Distribution of allelic fraction (AF) values across 80S and TOT mRNA fractions for the tranSNPs selected based on p < 0.05 and absolute value > 2SD. (C) Distribution of allelic fraction (AF) values across 80S and POL mRNA fractions for the tranSNPs selected based on p < 0.05. (D) Distribution of allelic fraction (AF) values across 80S and POL mRNA fractions for the tranSNPs selected based on p < 0.05 and absolute value > 2SD. (E) Heatmap plotting the delta allelic fraction (dAF) values for significant tranSNPs both in the POL-80S and 80S-TOT fractions. The trends of the corresponding POL-TOT values were also plotted.

### Haplotype analysis supports the identification of instances of allelic imbalance in monosome-seq

The RNA sequencing depth of our experiments allowed us to study over 5K heterozygous SNPs in the 80S monosome sequencing. Hence, we explored the consistency of allelic imbalance quantifications, exploiting haplotype phase reconstruction. For this analysis, we used the lists of putative tranSNPs (411 for the POL vs TOT, 359 for 80S vs POL, 200 for 80S vs TOT, and 358 for Nutlin POL vs TOT). We retrieved haplotypes consisting of at least three SNPs. Consistent with the proposal that the monosome represents the least translating RNA fraction we studied, the 80S vs. polysome comparison yielded the highest number of haplotypes (9, comprising 41 SNPs; **Figure 2A**). However, six of those nine haplotypes were in noncoding RNAs or pseudogenes. Comparing delta AF values within those haplotypes and considering their phase inferred from our RNA-sequencing data (**Table S2**), we observed high consistency in the directions of the imbalance data (**Figure 2B**). Information was retrieved for nine haplotypes across seven genes, totaling 26 SNPs, all of which were cis haplotypes (one haplotype with all reference alleles, the other with all alternative alleles). We then extended the analysis to genes with two tranSNPs identified (**Figure 2C**; SNPs marked by an asterisk). A total of 77 haplotypes were identified, of which 20 genes in the 80S vs POL comparison (corresponding to about 3.5% of analyzable haplotypes comprising two SNPs) (**Figure 2D**). Phasing data were inferred from our RNA-sequencing data for nine pairs, of which eight were *in cis* and one, ASB6, *in trans*, showing consistency with the dAF measurements (**Figure 2D**) again.

**Figure 2.**
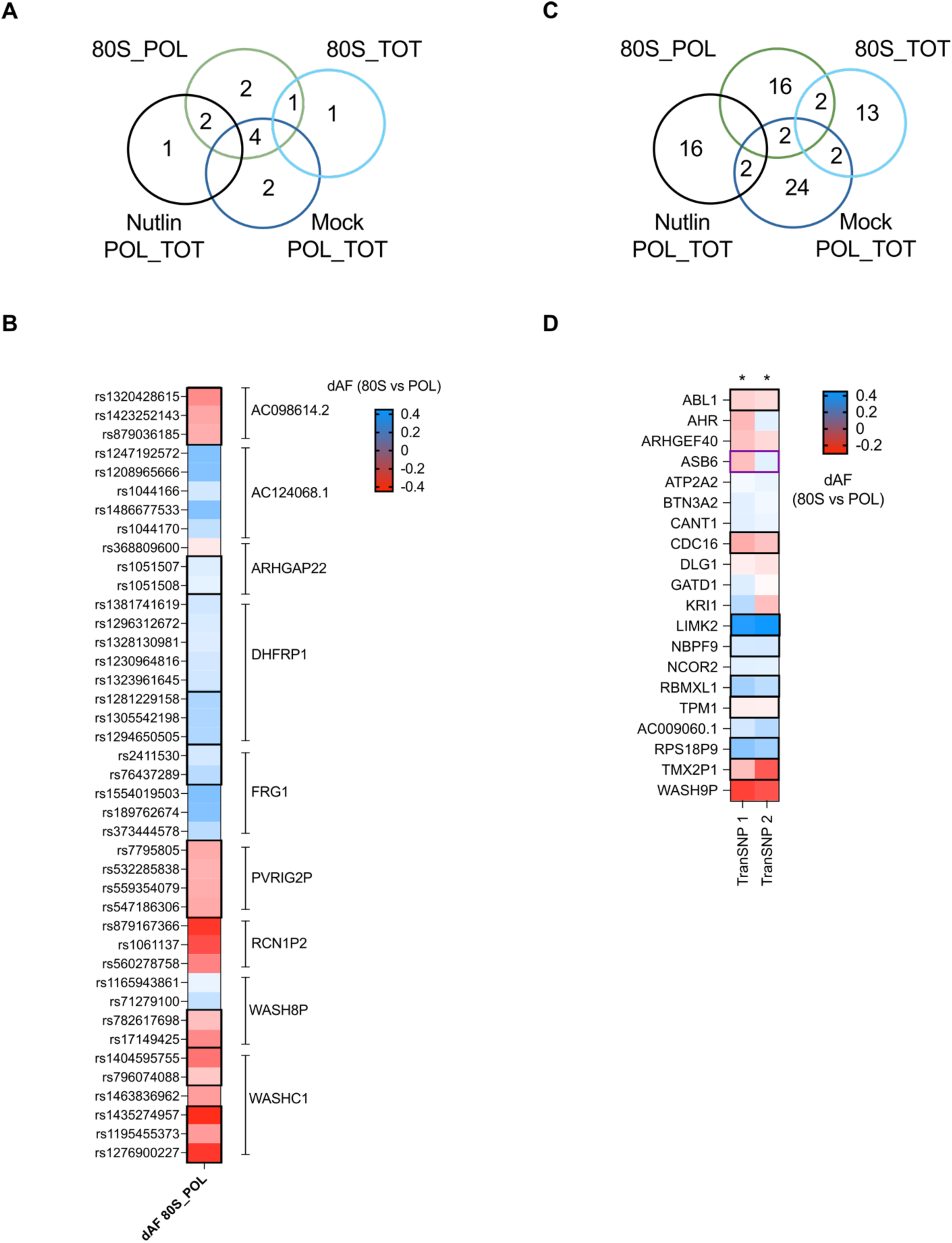
TranSNP haplotype analysis reveals consistent changes in allelic fractions. **A**) Venn diagram presenting the number of tranSNP haplotypes comprising three or more p-value <0.05 SNPs. **B)** Heatmap view of the dAF values for the nine haplotypes identified as imbalanced in the comparison between 80S monosome and polysome-seq. Six of those genes are either noncoding RNA or processed pseudogene. Haplotypes for which phasing data could be retrieved from RNA-sequencing data are highlighted by thicker boxes; two haplotype blocks were found for DHFRP1 and WASHC1 that could not be joined. **C**) Same as in A but focused on tranSNP haplotypes comprising two SNPs. **D**) Heatmap view of the dAF values for the 16 coding and four noncoding or pseudogene haplotypes identified as imbalanced in the comparison between 80S monosome and polysome-seq. Haplotypes for which phasing data could be retrieved from RNA-sequencing data are highlighted by thicker black (in cis) or purple (in trans) boxes. rsID codes are provided in Table S2.

Haplotype data were also inspected for the 17 cases retrieved from the 80S vs TOT comparison (**Figure S4**), although phasing data were retrievable only for three of them (CENPO, CNN2, UNC119B). Finally, we retrieved dAF data for all analyzable SNPs in the 80S haplotypes to explore whether consistent trends in dAF could also be observed for SNPs that did not show a statistically significant imbalance (**Figure S4**). However, long-read sequencing data would be required to conclusively validate these trends with phased haplotypes.

We concluded that haplotype analysis can support the identification of instances of allelic imbalance, extending the catalog of functional tranSNPs.

### UTR RPE-1 tranSNPs were validated by luciferase reporter assays

We performed luciferase reporter assays across 5’ and 3’ UTR tranSNP candidates to validate allele-specific translation efficiencies. We shortlisted candidates based on the magnitude of delta allelic imbalance, frequency of the alleles in the population, relevance of the host gene, and concordance of the imbalance with other SNPs *in cis* in the same mRNA, as inferred from haplotype analysis.

For the 14 chosen candidates, we cloned 109-nucleotide-long fragments of 5’ or 3’ UTRs containing either the reference or the alternative allele upstream or downstream of a Firefly luciferase reporter plasmid. We performed dual luciferase assays, using the Renilla luciferase as a normalizer of the Firefly luciferase signal first, and then Firefly mRNA levels to further correct the Firefly/Renilla ratios and focus on the post-transcriptional effect of the allelic variants.

Besides RPE-1 cells, in which this set of tranSNPs was identified, we extended the validation to HCT116 cells, as cancer cells are known to have rewired translation machineries (Fernández-Calero et al. 2020). **Figures 3A and 3B** show that most of the 5’UTR candidates did not exhibit significantly different translation efficiencies in RPE-1 cells, whereas significant differences were observed in HCT116 cells using this assay. Instead, we observed significant allele-specific differences among the 3’UTR candidates in both HCT116 and RPE-1 cells (**Figure 3C, 3D**). Notably, we detected a few cases in which the direction of allelic imbalance was opposite in RPE-1 and HCT116 cells, possibly due to differences in the availability or interaction with trans-factors, such as RNA-binding proteins (RBPs), miRNAs, or tRNAs.

**Figure 3.**
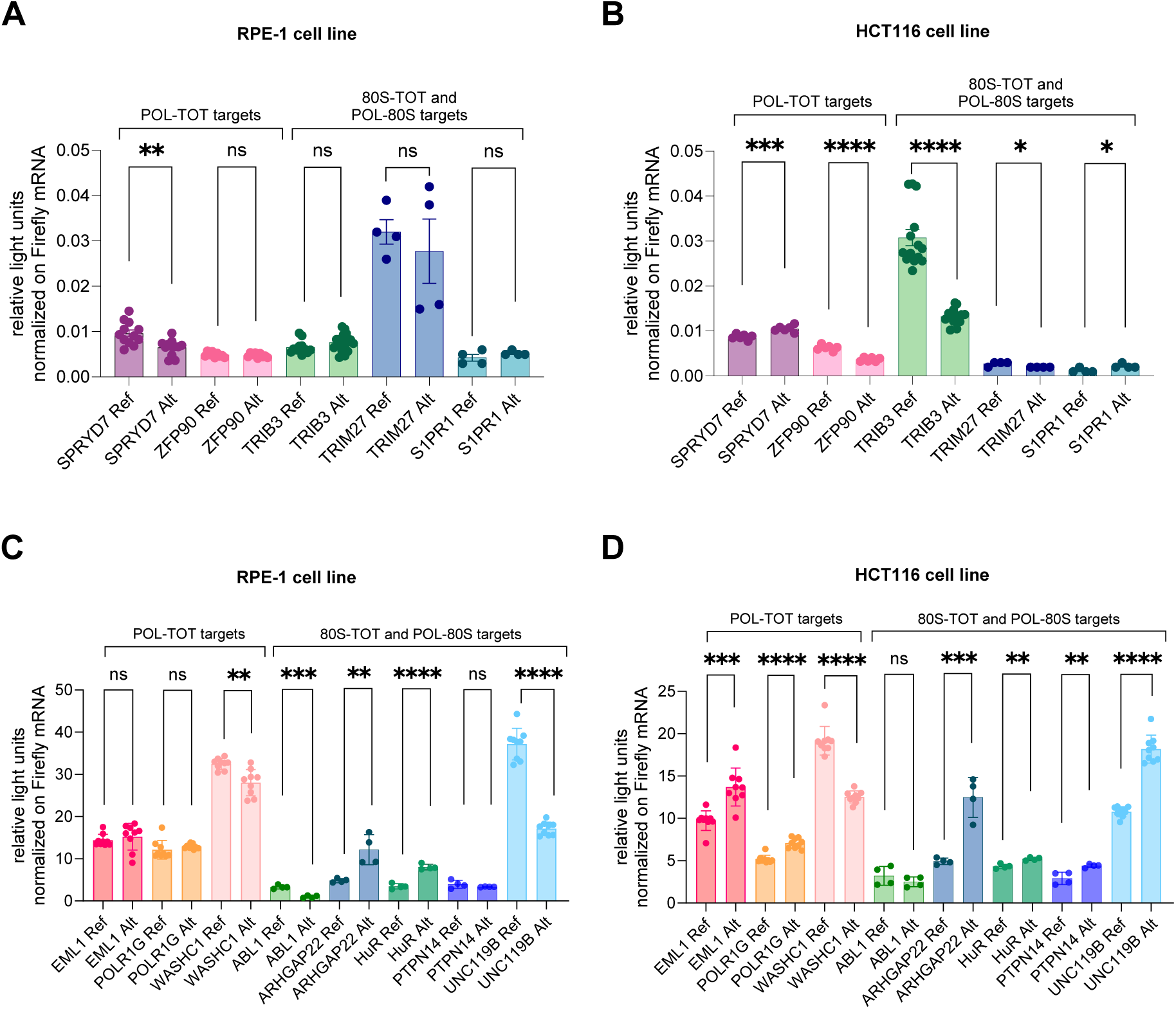
Allele-specific translation efficiencies for UTR RPE-1 tranSNPs were validated by luciferase reporter assays across RPE-1 cells and HCT116 cells. **A**) Luciferase reporter assay of 5’UTR constructs encompassing candidate tranSNPs in RPE-1 cells. ns, not significant, **p-value < 0.01, unpaired t-test. **B**) Luciferase reporter assay of 5’UTR constructs encompassing candidate tranSNPs in HCT116 cells. *p-value < 0.05, ***p-value < 0.001, ****p-value < 0.0001, unpaired t-test. **C)** Luciferase reporter assay of 3’UTR constructs encompassing candidate tranSNPs in RPE-1 cells. **p-value < 0.01, ***p-value < 0.001, ****p-value < 0.0001, unpaired t-test. **D**) Luciferase reporter assay of 3’UTR constructs encompassing candidate tranSNPs in HCT116 cells. **p-value < 0.01, ***p-value < 0.001, ****p-value < 0.0001, unpaired t-test.

### Coding RPE-1 tranSNPs influence ribosome stalling

We also selected four RPE-1 candidate tranSNPs located in the coding sequence (CDS) under the hypothesis that missense or silent changes in the CDS could influence ribosome sliding and stalling on mRNAs and, thus, translational pausing (Tsai et al. 2014). Similarly, candidates were selected based on the magnitude of allelic fraction imbalance, allele frequency in the population, RNA-sequencing coverage, host gene function, and, when available, concordance with other tranSNPs within the same mRNA, informed by haplotype analysis. (**Figure 4A**, **Table S3**).

**Figure 4.**
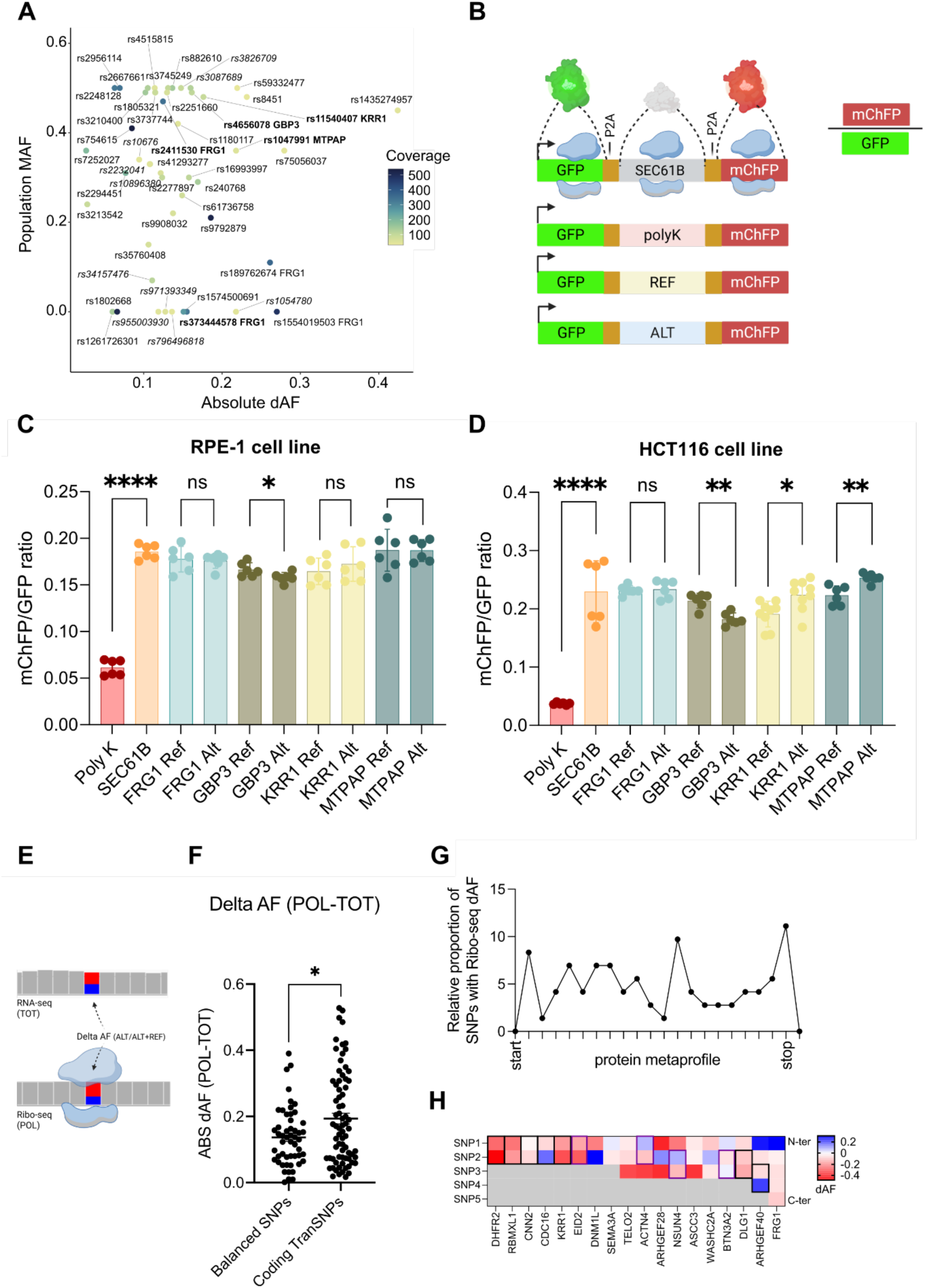
Coding RPE-1 tranSNPs influence ribosome stalling. **A)** Bubble plot representing the correlation between absolute delta allelic fraction (dAF), global population Minor Allelic Frequency (MAF), and coverage to guide the prioritization of coding RPE-1 tranSNPs. The highest absolute dAF value across conditions was plotted for each tranSNP. The selected tranSNPs for the stalling assay are shown in bold; the tranSNPs excluded from further analyses are shown in italics. **B)** Schematic representation of the ribosome stalling construct. The cloned sequences of interest (ALT/REF) were flanked by P2A sites to allow ribosome skipping and the cleavage of the three peptides separately from a single mRNA. The SEC61B sequence in the ribosome-stalling construct was used as a positive control. The ratio of mCherry to GFP signal intensities provides a measure of ribosome-scanning efficiency along the sequence of interest. **C)** mChFP/GFP ratios for the four selected tranSNPs measured in RPE-1 cells. **D)** mChFP/GFP ratios for the four selected tranSNPs measured in HCT116 cells. The fluorescence values were obtained using the Operetta High Content System. Data represent the mean ± SD of at least two independent experiments. ns, not significant, *p-value < 0.05, **p-value < 0.01, ****p-value < 0.0001, unpaired t-test. **E**) Scheme of the approach used to assess publicly available Ribo-seq data to quantify allelic imbalance on ribosome footprints. An IGV snapshot is used to visualize how the delta allelic fraction is calculated. The matched RNA-seq data is used as total (TOT) RNA coverage, while the Ribo-seq data is used as polysomal (POL) RNA. **F**) Comparison of Delta allelic fractions for the control and tranSNP groups. Ns = not significant; **p-value < 0.01, Ordinary one-way ANOVA with Tukey’s multiple comparisons test. **G)** Metaprofile of the distribution of 73 coding tranSNPs in 41 proteins from RPE-1 datasets (PRJNA281177 and PRJNA528026) (Table S2). Ticks in the x-axis correspond to 5% increments in relative protein length. **H**) Delta Allelic fractions calculated from RNA-seq and Ribo-seq data for the 18 proteins for which haplotype data consisting of at least two SNPs were retrievable. The data are ordered on the y-axis by haplotype length, while on the x-axis, SNPs are positioned relative to their positions in the primary sequence. Haplotypes for which phasing data could be retrieved from our RNA-sequencing data are highlighted in thicker black boxes when the reference and alternative alleles are in cis, and in purple when in trans.

For the four candidates, located in the FRG1, KRR1, GBP3 and MTPAP genes respectively, we cloned 109-nucleotide-long fragments of CDS, coding for 33 amino acids and containing either the reference or the alternative SNP allele in ribosome stalling plasmids, which consist of dual fluorescence plasmids characterized by the sequence of interest cloned between a GFP and an mCherry and flanked by two P2A sequences (Kriachkov et al. 2023) (**Figure 4B**). Impaired ribosome sliding by the sequence of interest is expected to reduce the mCherry signal, which lies downstream of the cloned sequence and thus the overall mChFP/GFP ratio. The SEC61B sequence, which is known to allow efficient readthrough, was used as a negative control for stalling. A positive control for stalling, containing a sequence coding for a stretch of 17 consecutive lysine residues (polyK), was also used. The ribosome stalling induced by all the cloned sequences was not severe and was comparable to that of the negative control; however, **Figures 4C** and **4D** showed allele-specific differences in ribosome stalling induced by tranSNPs, with a stronger effect observed in HCT116.

We also analyzed two publicly available RNA-seq data coupled with Ribo-seq in RPE-1 (BioProjects PRJNA281177 and PRJNA528026) (Tanenbaum et al. 2015) and used them as proxies for total RNA and polysomal RNA, respectively, to independently validate the imbalances observed with our method (**Figure 4E**). Data were obtained for 73 tranSNPs (**Table S3)**. Results were compared with a control group of 52 RPE-1 SNPs showing balanced AF in our datasets (**Table S3**). A significantly higher dAF was observed for the tranSNP group (**Figure 4F**), while no differences in the number of RNA-seq or Ribo-seq reads were noted (**Figure S4C, D**). We attempted to determine whether allelic imbalance in Ribo-seq data is associated with mRNA topography. Overall, there was no clear positional bias for the tranSNPs that could be analyzed by Ribo-seq, except for a slight enrichment towards the C-ter (**Figure 4G**). Next, we looked more specifically at instances of tranSNP-containing haplotypes. Ribo-seq data were available for eighteen cases (**Table S2**). We hypothesized that if a single SNP in a haplotype causes translation pausing or stalling, the direction of delta allelic fractions may be complementary between preceding and following SNPs. Phasing data were retrieved for eleven SNP pairs, of which seven were *in cis*. For three of them (CDC16, EID2, ARHGEF40), the haplotype dAF data support this conjecture (**Figure 4H**) (**Table S2**).

### The effect of missense tranSNPs can be explored by allele-specific proteomics

For coding tranSNPs that cause missense changes, it is, in principle, possible to explore allele-specific differences at endogenous levels by leveraging allele-specific proteomics. We recently validated this approach using the EIF4H and rs1554710467 SNV as an example (Hamadou et al. 2026). In the RPE-1 dataset, 40 missense tranSNPs in 35 proteins were found (**Table S3**). Compared to standard protein identification by mass-spectrometry, a specific peptide containing the SNP allele has to be detected. Furthermore, an added complexity arises from potential differences in the ionization efficiency of the two peptides corresponding to the tranSNP alleles, and, in the case of amino acid changes affecting arginine residues, also allele-specific differences in peptide length upon trypsin digestion. Indeed, of the 40 missense changes, 12 involved arginine residues in either the reference or the alternative alleles (**Table S3**). However, Lys-C, Glu-C, and chymotrypsin were less efficient than trypsin. Using label-free shotgun proteomics, peptides from 17 of the 35 proteins were identified (**Figure 5A**). However, only in 4 cases a peptide spanning the biallelic position was detected (**Figure 5B**), and in the case of FRG1, only the reference peptide was found. In general, transcripts and protein relative abundance were correlated, but abundance alone did not predict the likelihood of finding the desired peptide for allelic quantification (**Figure 5B**). To enrich for the DLG1, CAST, and CHTF18 proteins, total extracts were resolved by SDS polyacrylamide gel electrophoresis, stained with Coomassie, and gel bands were excised, cut, destained, and in-gel digested using trypsin (see Methods). A targeted approach was then chosen to obtain relative quantifications of the reference and alternative peptides (**Figure 5C**).

**Figure 5.**
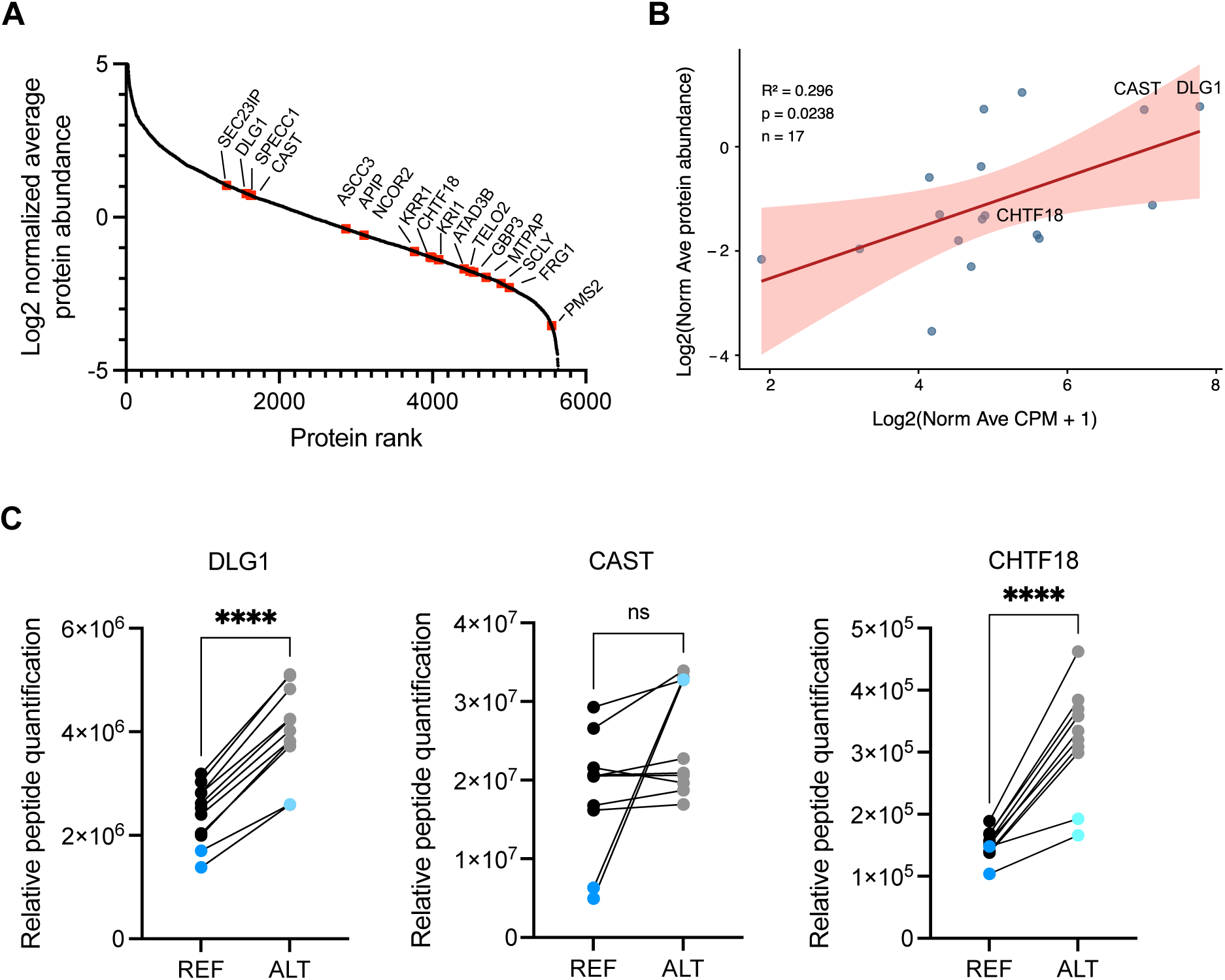
Exploration of missense tranSNPs by label-free proteomics. **A**) Whole proteome using label-free shotgun analysis. Relative protein abundance was ranked based on the aggregated results of three biological replicates, each analyzed by three injections. The ranked positions of proteins from the list of missense transSNPs are highlighted in red. **B**) Comparison between mRNA expression (CPM, average of the biological replicates) and relative protein abundance from RPE-1 whole proteome analysis. Although 17 proteins with missense tranSNPs were identified, in only three cases, highlighted by the protein name, peptides comprising the biallelic variant were detected by mass spectrometry. **C**) Relative quantification of the reference (REF) and alternative (ALT) peptides for DLG1, CAST, and CHTF18 by label-free targeted analysis, based on three biological replicates. The two data points in blue refer to analyses of technical replicates of a whole protein lysate sample, while those in black refer to three biological samples prepared by in-gel digestion, each tested with two technical replicates. ns, not significant; **** p< 0.0001, two-tailed paired t-test.

### TranSNPs are associated with allele-dependent RNA-binding protein interactions and clinical outcomes

To begin exploring trans-factors potentially involved in allele-specific interactions by targeting tranSNP-containing sequences, binding predictions were generated using the TESS software (Schug 2008), starting with a list of 53 RNA-binding proteins (RBPs) with a consensus motif (Cook et al. 2011). These were used to compute binding scores for the reference and alternative alleles for 280 tranSNPs (**Table S1**). RBPs motifs overlapping a SNP site were found for 194 of the 280 cases (**Table S4a**). This list was further refined by filtering for a binding score of at least 75% of the maximum score for one of the two alleles and considering the magnitude of the impact of each SNP allele to result in at least a 50% change in the binding score prediction. This conservative approach led to a shortlist of 110 among the 280 tranSNPs with at least one prediction of allele-prevalent binding by RBPs. A total of 137 allele-prevalent binding predictions were obtained (**Table S4b**) for 11 RBPs. About 65% of the predicted sites mapped to exonic positions. Focusing on the 17 tranSNPs for which validation reporter assays were developed, predictions of allele-specific binding were obtained for 7 SNP sites and five different RBPs (**Table 1**).

**Table 1.** Predictions of allele-specific or allele-prevalent binding were obtained for 7 SNP sites and five different RBPs.

| rsid | gene | RBP motif | refScore | altScore | maxScore | % of change |
| --- | --- | --- | --- | --- | --- | --- |
| rs140546414 | ABL1 | RBMX | 0.13 | 5.17 | 5.86 | 86.02 |
| rs12983784 | ELAVL1 | SFRS1 | 1.87 | 4.19 | 4.62 | 50.26 |
| rs12983784 | ELAVL1 | EIF4B | 3.25 | 8.98 | 10.49 | 54.61 |
| rs1357255885 | PTPN14 | EIF4B | 2.22 | 8.88 | 9.98 | 66.73 |
| rs1357255885 | PTPN14 | RBMX | 5.86 | 0.82 | 5.86 | -86.02 |
| rs1063181 | SPRYD7 | SFRS1 | 3.57 | 1.25 | 4.62 | -50.26 |
| rs1063181 | SPRYD7 | EIF4B | 5.75 | None | 7.08 | -81.14 |
| rs1404595755 | WASHC1 | FUS | 0.60 | 7.89 | 7.89 | 92.44 |
| rs1404595755 | WASHC1 | SFRS1 | 3.57 | 1.25 | 4.62 | -50.26 |
| rs3737577 | S1PR1 | ELAVL1 | 2.08 | 4.40 | 4.40 | 52.73 |
| rs4656078 | GBP3 | RBMX | 4.91 | 0.52 | 5.86 | -74.90 |

As in our previous studies (Valentini et al. 2021; Hamadou et al. 2025; Valentini et al. 2022), we interrogated TCGA data (Liu et al. 2018) using germline genetic information to explore the potential clinical relevance of the identified tranSNPs. Among the 7 tranSNPs validated by reporter assays and for which allele-specific RBP binding was predicted, four were associated with clinical outcomes under either a recessive or a dominant model for the alternative allele across OS, DSS, DFI, and PFI endpoints (**Table S5**). Three of them, rs12983784 in the ELAVL1 3’UTR, rs3737577 in the S1PR1 5’UTR, and rs1063181 in the SPRYD7 5’UTR, had prognostic value in ovarian cancer (**Figure 6A-C**), suggesting a potential functional link between RNA-protein interactions and clinical phenotypes.

**Figure 6.**
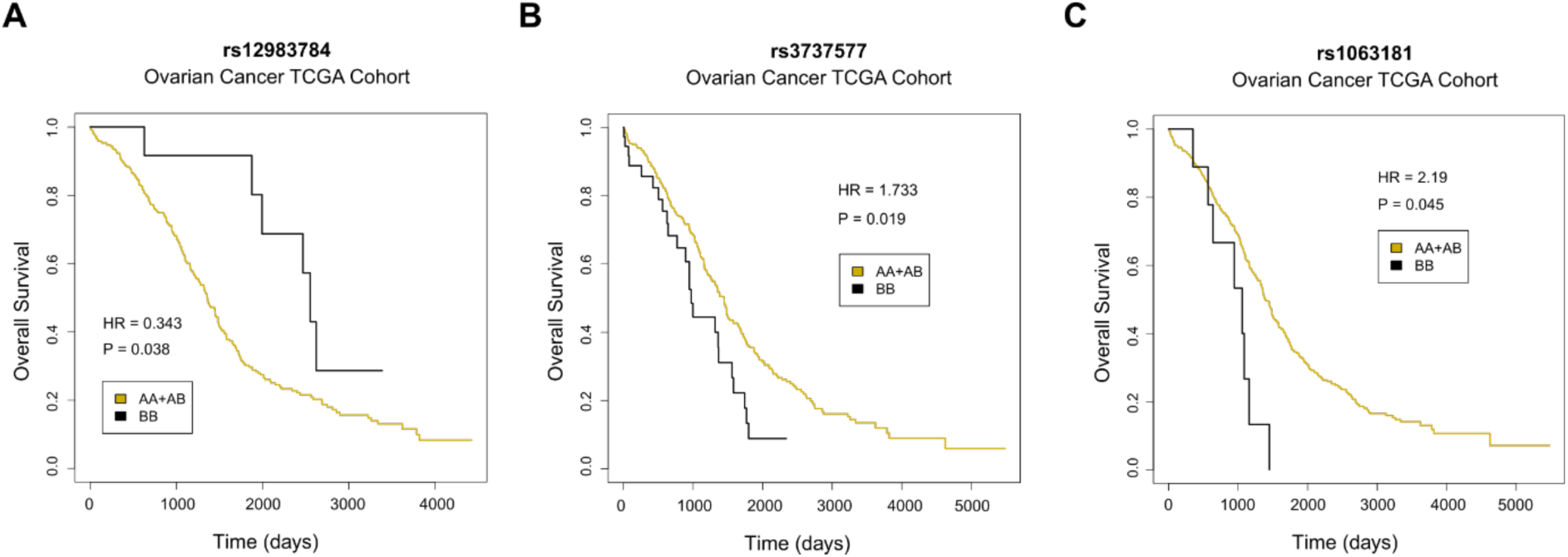
TranSNPs can be correlated with distinct clinical outcomes in TCGA. The rs12983784 **A**), rs3737577 **B**), and rs1063181 **C**) genotypes stratify ovarian cancer patients across a TCGA cohort for the Overall Survival endpoint. Kaplan-Meier curves plotting the Overall Survival under a recessive model (B = ALT allele) are presented along with the hazardous ratio (HR) and the p-values.

## Discussion

Our study establishes that allele-associated differences in post-transcriptional mRNA regulation can also be identified and studied in non-transformed cells. The dataset generated in RPE-1 cells showed that the 80S-bound mRNAs represent the least translated fraction in RPE-1 and improve the resolution in the identification of tranSNPs. However, luciferase reporter assays performed in parallel in RPE-1 and HCT116 cells to validate UTR tranSNPs revealed cell-line-specific differences in the magnitude and direction of the imbalance between reference and alternative alleles, especially among the 3’ UTR targets arising from the comparison with 80S-bound mRNAs, suggesting a potential different use of the 80S monosome (Figure 3). This finding is not entirely unexpected, as the translational potential of the 80S monosome remains debated (Heyer and Moore 2016; Schneider et al. 2022), and recent evidence in Drosophila melanogaster indicates that it can be highly tissue-dependent (Blandy et al. 2025). This feature would limit the possibility of generalizing the use of 80S-bound mRNAs as a proxy for untranslated mRNAs. Interestingly, in Drosophila, 80S monosomes appeared to be translationally active during developmental stages or in organs where translation regulation requires more precise control, involving structured 5’UTRs or upstream ORFs. Conversely, 80S monosomes appeared inactive in the testis and ovary, where polysomes were the major sites of protein synthesis (Blandy et al. 2025). Oncogenic signaling, or, conversely, nutrient starvation or hypoxia, can impact both global and mRNA-specific translation (Lee et al. 2020; Mossmann et al. 2018; Koritzinsky et al. 2006). Hence, specific conditions or perturbations may help predict the translational potential of the 80S and its value for pursuing the genetic bases of allelic differences in translational regulation.

Moreover, while we observed that the comparison between polysome and 80S-bound mRNAs provides the highest resolution in tranSNPs identification in RPE-1 cells, we further found that comparing 80S with total cytoplasmic mRNAs has higher discriminatory potential, consisting in larger delta allelic fractions, than comparing polysomal with total mRNAs (Figure 1E), suggesting that the polysomal presence in the total mRNA fraction contributes more than the 80S in the total cytoplasmic mRNA fraction in this cell line.

We used luciferase-based assays to evaluate allele-specific differences in translation efficiency for 5’ and 3’ UTR targets. Differences in the magnitude of the effect of the cloned fragments were observed between 5’ UTR and 3’ UTR constructs across both RPE-1 and HCT116 cells. These can be attributable to the use of two different plasmid scaffolds that contain a distinct constitutive promoter and may differ in the translation efficiency of the reporter: pGL3p plasmid was used for cloning the 5’ UTR targets, and pGL4.13, harboring a codon-optimized Firefly luciferase, for cloning the 3’ UTR targets. Both types of plasmids express high levels of the reporter, which could reduce the assay’s dynamic range; however, we adopted double normalization of the light units, using both the control Renilla luciferase activity and the relative levels of firefly mRNA measured by qPCR in the same extracts used for the luminescence assay. Importantly, in RPE-1 cells tranSNPs selected exploiting the 80S monosome-seq were validated more efficiently. As noted, comparing results in RPE-1 and HCT116 showed examples of striking cell line differences in allelic impact, suggesting strong context-dependent effects possibly related to the different use of the 80S monosome or the relative expression of trans factors.

Tissue-specific differences in the expression of RNA-binding proteins and microRNAs, tRNAs, epitranscriptomic writers and readers can underlie differences in the kinetics or extent of pre-mRNA splicing, mRNA export, subcellular localization, poly-adenylation, translation initiation and elongation, all features that can also impact the potential to identify distinct flavors of biallelic mRNA fate (Koletsou and Huppertz 2025; Xiang et al. 2024; Luo et al. 2025; Fabbri et al. 2021; Tian et al. 2026). These post-transcriptional regulatory layers may lead to tranSNP positional biases along mRNA sequences in cases of heterozygous haplotypes, as we sought to explore in this study. High-depth RNA-sequencing, paired-end protocols, and especially long-read sequencing may enable the identification of tranSNPs at the transcript rather than the gene level in future studies.

While the analysis pipeline was specifically developed to prioritize functional SNPs in post-transcriptional gene regulation, establishing the causal effects of cis-alleles remains challenging due to low throughput and limitations of reporter assays that must account for post-transcriptional mRNA regulation. In this respect, fluorescence-based assays are, in principle, superior to luminescence-based assays, although relative sensitivity can be a limitation. Even more complex is the identification of causative trans-factors. For example, the validation of allele-specific binding of an RNA-binding protein to a tranSNP-containing site requires either experiments with allelic resolution in cells, such as a RIP protocol followed by bound RNA analysis, or edited isogenic cells homozygous for the two tranSNP genotypes, or else the comparison of binding in a collection of heterologous cells that already harbor distinct tranSNP genotypes, but may also differ for the relative expression of target RBPs. In this respect, the data we presented on predicted tranSNP-altered RNA-binding proteins’ binding sites represent an exploratory first step towards a mechanistic understanding. The same can be said for the prognostic features in TCGA cancer patient cohorts stratified based on tranSNP genotype. These limitations notwithstanding, the fact that among the 17 candidate tranSNPs chosen for validation, for two of them allele-specific differences in reporter activity, RBP binding, and patient-survival were found suggests that our dataset can be a useful resource to discover functional genetic variants, particularly in the untranslated regions of mRNAs.

The translation efficiency of coding tranSNPs has been assessed instead by dual fluorescence assays designed to detect instances of ribosome stalling. We observed allele-specific differences in ribosome stalling capacity. The magnitude of the effect was expectedly modest compared to the positive control, which features a strong stalling sequence consisting of 17 consecutive lysine residues. We estimated that the magnitude of the effect of the single amino acid substitutions we studied is similar to the impact observed with 10 copies of a GR-coding hexanucleotide repeat derived from c9orf72 intron 1 (Kriachkov et al. 2023). We also measured allelic imbalance of a number of tranSNPs using publicly available Ribo-seq and RNA-seq datasets. However, it has to be taken into consideration that Ribo-seq footprints do not distinguish monosomes from polysomes, although the positional resolution, coupled to haplotype phasing when multiple heterozygous SNPs are present, could help exploring if allele-specific effects are related to differences in translation initiation, or in ribosome-pausing at specific sequence contexts, or to reduced availability of one allelic transcript to the translation machinery caused for example by mRNA compartmentalization. In this latter case, delta allelic fractions consistent in direction along the mRNA sequence are expected, while in the former two cases, a change in the direction of the delta allelic fractions along the mRNA sequence could be anticipated. In fact, if a tranSNP represents a ribosome pause site, ribosomes might queue before it, but not after (Figure 4H). While limited by sequencing depth and resulting coverage at SNP sites, our results suggest that Ribo-seq data, coupled with biallelic markers in the coding sequence, could reveal genetic variants affecting subtle mRNA translation features. It would be interesting to explore if variations to the Ribo-seq protocol, such as polysome-seq (Floor and Doudna 2016; Yoshikawa et al. 2018) or Ribolace (Clamer et al. 2018), i.e., methods that enable the isolation of translating ribosomes, could increase the allelic resolution at tranSNP sites. The isolation of specific polysome species using optimized sucrose gradients (Liang et al. 2018) represents further improvements to the experimental pipeline. In addition to 80S monosome sequencing, the fractions lighter than ribosomal subunits and the 40S fraction could be used as a source of cytoplasmic mRNA not undergoing translation. However, considering the typical distribution of relative mRNA abundance along the sucrose-gradient profile (Panda et al. 2017; Hamadou et al. 2025), mRNA species in those fractions may be under-represented. Finally, nuclear RNA could also be considered to collectively enable capturing distinct aspects of post-transcriptional mRNA regulation, empowered by the biallelic marker resolution.

TranSNPs that result in missense changes are potentially amenable to validation at the endogenous level in heterozygous cells using allele-specific proteomics. We recently validated this approach using a missense SNV in EIF4H (Hamadou et al. 2026). Here, we explored whether a simplified mass spectrometry approach that did not include isotype-labeled synthetic peptide spike-ins for each chosen target could still enable tranSNP validation. Results confirmed that MS sensitivity remains suboptimal, especially given the need to quantify a specific, biallelic peptide and the complexity introduced by missense changes affecting arginine or methionine residues, which can alter peptide properties, such as ionization or oxidation. In fact, starting from 35 candidate proteins, only in three cases could both the reference and alternative peptide relative quantifications be obtained, and a significant imbalance was observed in two of them (Figure 5). For the CAST target, the difference in the relative quantification of the reference peptide between whole lysates and SDS-PAGE and in-gel digestion remains unexplained.

Nutlin treatment in RPE-1 cells led to broad changes in the transcriptome consistent with p53 activation and cell cycle arrest (Tiu et al. 2021; Andrysik et al. 2017; Rizzotto et al. 2020). The treatment-dependent changes in gene expression were largely coordinated at the transcriptional and translational levels, although a trend toward high-magnitude repression was apparent in the translatome (Figure S2). Nutlin treatment was shown to alter global translation control by modulating p53-dependent EIF4E-BP1 and ribosome biogenesis factors, RNA-binding proteins, and microRNAs (Lindström et al. 2022; Marcel et al. 2013; Tiu et al. 2021; Marcel et al. 2018; Hermeking 2012). However, Nutlin treatment of RPE-1 cells did not result in a global change in the detection of tranSNPs. Hence, our analysis pipeline can be applied in a broader context, beyond cancer research, as a tool to map genetically driven inter-individual sources of phenotypic diversity, while providing clues to guide mechanistic studies.

## Methods

### Cell culture

The human non-cancer cell line of retinal pigmented epithelia RPE-1 was cultured in Dulbecco’s Modified Eagle Medium and Ham’s F12 (DMEM/F12) supplemented with 10% fetal bovine serum (FBS), 100 units/ml penicillin, 100 mg/ml streptomycin antibiotic mix (GIBCO), and 1% L-Glutamine (GIBCO). The cells were maintained in a humidified atmosphere at 37°C and 5% CO2. Cells were split twice a week.

### RNA extraction

RNA was extracted using the NucleoSpin® RNA extraction Kit (Macherey-Nagel) according to the manufacturer’s instructions. RNA purity and concentration were assessed by NanoDrop™ Spectrophotometer (Thermo Fisher Scientific).

### cDNA and RT-qPCR

RNA was retrotranscribed using the RevertAid First Strand cDNA Synthesis Kit (Thermo Fisher Scientific) and random hexamer primers according to the manufacturer’s instructions. One µg of RNA was retrotranscribed and the cDNA was diluted 1:8 in water, equivalent to a final concentration of 6.25 ng/µl. RT-qPCR was performed using the PowerUp SYBR mix (Applied Biosystems) in 10 µl, using 12.5 ng of cDNA as input and 0.4 µM of each primer. Primers were designed using Primer Blast (NCBI) and are listed in **Table S6**. The QuantStudio5 instrument was used to amplify the cDNA and measure gene expression. Technical triplicates were performed, and average Cq values were normalized to housekeeping genes (dCt) and to controls (ddCt) when possible.

### Polysome profiling and RNA sequencing

Cells were grown in 150 mm dishes to 70-80% confluence and incubated for 10 min with 100 µg/ml cycloheximide at 37°C to block protein synthesis and immobilize ribosomes on mRNAs. Cells were then rinsed with ice-cold PBS 1X containing 50 µg/ml (or 100 µg/ml) cycloheximide, scraped, and pelleted. The cell pellet was lysed with 600 µl of ice-cold lysis buffer (20 mM Tris-HCl pH 7.5, 130 mM KCl, 30 mM MgCl2, 2.5 mM DTT, 0.2 % NP-40, 0.5 % sodium deoxycholate, 0.2 mg/ml heparin, 0.2 U/µl RNasin® Ribonuclease Inhibitor (Promega), cOmplete™ Mini Protease Inhibitor Cocktail 1X (Roche), 100 µg/ml cycloheximide) by incubation on ice for 10 minutes and centrifugation at 12000 g at 4 °C for 10 min to remove cellular debris and nuclei. The cytoplasmic lysate was then loaded on a 15-50% linear sucrose gradient dissolved in salt buffer (100 mM NaCl, 20 mM Tris-HCl, pH 7.5, and 5 mM MgCl2) and processed by ultracentrifugation in a Beckman SW41 rotor at 40000 rpm for 1 hour and 40 minutes at 4°C. The gradient was then fractionated in 1 ml fractions, and the absorbance was measured at 254 nm by the Teledyne Isco model 160 gradient analyzer equipped with a UA-6 UV/VIS detector. RNA was extracted from the fractions using TRIzol® based on the manufacturer’s instructions. RNA-seq libraries were prepared using the Illumina Stranded mRNA kit and sequenced at the CIBIO NGS Core facility using an Illumina NovaSeq.

### RNA-seq data processing, differential expression analysis and tranSNP identification

Paired-end RNA-seq data from total, 80S/monosomal and polysome-associated RNA fractions were processed from raw FASTQ files to generate BAM files for gene-level expression analysis. Adapter sequences and low-quality bases were removed using Trimmomatic (Bolger et al. 2014). Trimmed reads were aligned to the human reference genome GRCh38 using STAR (Dobin et al. 2013), generating one BAM file for each sample. Gene-level expression was quantified by counting reads mapping to genes annotated in the reference genome using the Rsubread R package (Liao et al. 2019). Differential gene expression analysis was performed with the edgeR R package (Love et al. 2014) to identify genes showing significant expression changes across RNA fractions. Pairwise comparisons were performed among total, 80S/monosomal and polysome-associated RNA fractions. Genes were considered differentially expressed based on statistical significance, defined as adjusted p-value < 0.01, effect size, defined as absolute log2 fold change > 1, and average expression signal, defined as logCPM > 0. Functional enrichment analysis was performed separately on the lists of up-regulated and down-regulated genes obtained from each pairwise comparison using the Metascape web portal (Zhou et al. 2019), which integrates Gene Ontology terms, pathway databases, and additional functional annotations to identify significantly over-represented biological processes and molecular pathways. For tranSNP identification, the aligned BAM files were subjected to additional post-alignment processing to improve the reliability of allele-specific read counts at SNP positions. PCR duplicates were removed using the Picard MarkDuplicates module. The resulting BAM files were then processed with GATK to perform RNA-seq-specific refinement steps, including splitting of reads spanning splice junctions, local realignment, and base-quality recalibration. These additional processing steps were applied specifically for allelic fraction estimation and were not used for differential gene expression analysis.

TranSNP identification was performed following the workflow previously developed for the detection of single-nucleotide variants associated with allele-specific translation potential (Valentini et al. 2021) and implemented as described in Hamadou, Alunno et al (Hamadou et al. 2025). Briefly, post-processed BAM files were interrogated at dbSNP-annotated SNP positions located in exonic and untranslated regions. Pileup at SNP positions was performed using ASEQ with parameters mode=0, mbq=20 and mrq=20. Allelic fractions were calculated as the number of reads supporting the alternative allele divided by the total number of reads supporting the alternative and reference alleles. Heterozygous SNPs with sufficient read coverage were retained, and allelic fractions were compared across total, 80S/monosomal and polysome-associated RNA fractions. Candidate tranSNPs were defined as variants showing consistent allelic imbalance between RNA fractions across biological replicates.

### Cloning of tranSNP-containing sequences for reporter assays

Forward and reverse primers containing the restriction sites and the sequence of interest (insert) were designed (sequences are reported in **Table S6**), resuspended to a final concentration of 100 µM, and simultaneously annealed and phosphorylated by mixing 1:1 (8.5 µl each) in a final volume of 20 µl with 2 µl of 10X DNA ligase buffer (Thermo Fisher Scientific) and 0.5 µl of PNK kinase (New England Biolabs). Annealed inserts were diluted 1:500 in water. In the meantime, backbone plasmids (about 1 µg) were double-digested using restriction enzymes for 3 hours at 37°C using 0.5 µl of each enzyme in a final volume of 20 µl. The digestion product was purified from gel using the Promega Gel Extraction and PCR Clean-Up Kit and quantified using the NanoDrop™ Spectrophotometer (Thermo Fisher Scientific). About 50 ng of digested backbone were incubated with 1-2 µl of annealed insert (or water in the insert negative sample) and 2 µl of DNA T4 Ligase (Thermo Fisher Scientific) in a final volume of 20 µl for 2 to 3 hours at 22°C. The ligation product was entirely transformed into competent DH5α E. Coli bacteria (see Bacteria Transformation Section), and all bacteria were plated onto Ampicillin Liquid Broth (LB) Agar plates and incubated overnight at 37°C. Single colonies were picked and cultured in 5 mL of liquid LB supplemented with Ampicillin overnight at 37°C in a shaking incubator. Plasmid DNA was extracted using the PureYield™ Plasmid Miniprep System (Promega), quantified, and the correct insertion was verified via diagnostic digestion and Sanger sequencing.

### Luciferase-based reporter assays

5000 cells were seeded in a 96-well plate format in 100 µl of medium. After 24 hours, cells were transfected with 75 ng of DNA from the cloned constructs on Firefly backbone plasmids (pGL3 promoter for 5’UTR targets and linker pGL4.13 for 3’UTR targets, see Table 6; a “linker” was added to the pGL4 plasmid to generate an EcoRI and an NdeI restriction site around the XbaI site), and 25 ng of DNA from the Renilla luciferase plasmid. Plasmid DNA was diluted in Opti-MEM™ Reduced Serum Medium (Thermo Fisher Scientific), and FuGENE® HD (Promega) was used as a transfection reagent in a 4:1 ratio. The transfection reaction was assembled in a 5 µl of volume and incubated at room temperature for 15 minutes before being added dropwise to the cells. After 48 hours from the transfection, cells were washed with 150 µl of PBS 1X and lysed using 50 µl per well of Passive Lysis Buffer 1X (PLB) in shaking at 500 rpm for 15 minutes. PLB and the following reagents were part of the Dual-Luciferase® Reporter™ Assay System. 10 µl of lysate were then transferred into white 384-well plates. 10 µl of Luciferase Assay Reagent (LAR), which is the Firefly Luciferase substrate, were added to each well. Upon brief shaking, the Firefly luciferase signal was detected using the Varioskan™ LUX Multimode Microplate Reader (Thermo Fisher Scientific). Three readings were performed. STOP&Glo reagent was then diluted 1:50 in the provided buffer, and 10 µl were added to each well. Upon brief shaking, the Renilla luciferase signal was detected using the same instrument; three more readings were performed. Data were analyzed by calculating the ratio between the Firefly and the Renilla luminescence signals for each well. The remaining, untransferred 40 µl of lysate were collected, and RNA was extracted using TRIzol™ (Invitrogen). RNA was then quantified, retrotranscribed, and Firefly mRNA was amplified via RT-qPCR (as in the cDNA and RT-qPCR Section).

### Ribosome stalling assay

109-nucleotide long sequences (coding for 33 amino acids) comprising either the alternative or the reference version of the CDS tranSNPs were cloned in dual reporter ribosome stalling plasmid constructs (Kriachkov et al. 2023) upstream of an mCherry and downstream of a GFP sequence, separated by P2A sequences which induce the ribosomes to skip the formation of peptide bonds without interrupting translation, allowing the translation of separate peptides from a single mRNA (Lin et al. 2013). Cells were seeded in a 96-well plate format and, after 24 hours, transfected with 100 ng of the above-mentioned ribosome stalling plasmids using the FuGene transfection reagent (Promega) in a 4:1 ratio. A known staller sequence (coding for 17 consecutive lysines, polyK) cloned in the same plasmid was used as a positive control. A SEC61B sequence (106 amino acids), which is known to allow read-through, was cloned into the same plasmid and used as a negative control. GFP and mCherry signals, and digital phase contrast were detected after 24-, 36-, and 48-hours post-transfection using the Operetta High Content Imaging System (Perkin Elmer). GFP and mCherry intensities were used to calculate mChFP/GFP ratios, which were used as proxies for ribosome stalling. Low mChFP/GFP ratios indicated that stalling events occurred.

### Proteomic analysis: sample preparation and selection of Peptides for Targeted Proteomics (PRM)

RPE-1 cells were harvested and processed in RIPA buffer for 30 min at 4 °C, followed by centrifugation to remove cellular debris. Protein concentration was quantified via the BCA assay. For each sample, 20 μg of protein underwent reduction with 10mM Tris (2-carboxyethyl) phosphine (TCEP) followed by reduction for 30 min at room temperature using 20 mM iodoacetamide (IAA) at 25 °C for 30 min in the dark. Protein digestion was conducted using carboxylate-modified magnetic beads (GE Healthcare) according to the SP3 protocol (Hughes et al. 2019). Briefly, washed beads were mixed with proteins at a 1:10 (w/w) ratio. Acetonitrile (ACN) was added to a final concentration of 70% (V/V), and the mixture was incubated at room temperature for 18 min. After magnetic separation and washing (twice with 70% ethanol, once with ACN), beads were resuspended in digestion buffer (50 mM NH4HCO3, 5 mM CaCl2) containing different proteases (Lys-C, Glu-C, ChymoTripsin or Trypsin) at a 1:15 enzyme-to-protein ratio. Following an overnight incubation at 37 °C, peptides were desalted using BioSPE® PurePep Tips (Affinisep), according to the manufacturer’s instructions. To enrich proteins of interest, lysates were separated via SDS-PAGE using Bolt 10% Bis-Tris minigels (Thermo Fisher Scientific) and visualized with Imperial Protein Stain (Thermo Fisher Scientific). Gel regions corresponding to 25–45 kDa and 70–100 kDa were excised and minced into ∼1 mm³ fragments. These pieces were destained using a 1:1 mixture of 100 mM NH4HCO3 and acetonitrile (ACN), followed by dehydration in 100% ACN and centrifugal evaporation (SpeedVac). Proteins were reduced with 10 mM DTT and alkylated with 55 mM IAA. Following washes with 100 NH4HCO3 and ACN, the dried gel pieces were saturated with trypsin (12.5 ng/μL) on ice for 45 min. Proteolysis was conducted overnight at 37 °C. Peptides were recovered through sequential extraction using 30% ACN/3% trifluoroacetic acid (TFA) and pure ACN. The pooled extracts were concentrated via SpeedVac, acidified to pH < 2.5 with 1% TFA, and desalted using laboratory-made C18 StageTips. Peptide samples were reconstituted in 0.1% formic acid and stored at −20 °C prior to shotgun and PRM analysis.

Selected proteins were subjected to in silico tryptic digestion to identify peptide sequences harboring the SNVs of interest. Because peptide selection was constrained by the specific biallelic variant sites, it was not always feasible to meet all standard criteria typically applied for the choice of optimal quantotypic peptides for PRM assays (Rauniyar 2015). In particular, the only sequence-related limitation was the unavoidable inclusion of cysteine residues requiring carbamidomethylation. For each candidate peptide, the most suitable precursor charge state was determined through manual evaluation of mass spectrometric performance, considering factors such as signal-to-noise ratio, transition intensity, chromatographic peak shape, and integrated peak area.

### Proteomic analysis: LC–MS/MS and Data analysis

For shotgun label-free quantification, digested peptide samples were separated using a Dionex 3000 nanoLC RSLC system (Thermo Fisher Scientific) coupled to a heated reversed-phase analytical column (75 µm internal diameter, 1.7 µm particle size, 100 Å pore size; CoAnn Technologies) maintained at 40 °C. Chromatographic separation was performed using a binary mobile phase system composed of 0.1% formic acid in water (buffer A) and 0.1% formic acid in acetonitrile (buffer B). Peptides were eluted at a flow rate of 300 nL/min using the following gradient: 5–25% buffer B over 80 min, 25–40% over 3 min, 40–98% over 2 min, followed by a 15-min wash at 98% buffer B. Eluting peptides were analyzed on an Orbitrap Fusion Tribrid mass spectrometer (Thermo Fisher Scientific) operating in data-dependent acquisition mode, with a spray voltage of 2.1 kV and an ion transfer tube temperature of 200 °C. Full MS scans were acquired in the Orbitrap at a resolution of 120,000 (FWHM at m/z 200), with an AGC target of 4 × 10⁶ and a maximum injection time of 246 ms. Survey scans covered a mass range of m/z 250–1500. Dynamic exclusion was enabled with a 40-s exclusion window. Each survey scan was followed by MS/MS fragmentation using higher-energy collisional dissociation (HCD) at 30% normalized collision energy within a 3 s duty cycle. MS/MS scans were acquired either in the Orbitrap or in the ion trap, with maximum injection times of 54 ms and 35 ms, respectively, and an AGC target of 2.5 × 10⁴.

For targeted PRM analysis, peptides containing the specific biallelic amino acid variants were monitored. Liquid chromatography conditions were identical to those used for shotgun analyses. Selected precursor ions were included in an isolation list containing peptide masses and charge states (**Table S7**). PRM scans were acquired in the Orbitrap at a resolution of 120,000, with an AGC target of 4.0 × 10⁵, a maximum injection time of 246 ms, and HCD fragmentation at 30% normalized collision energy. To minimize sample carryover and ensure consistent analytical performance, blank injections and quality control samples were analyzed between runs. Instrument performance was continuously monitored throughout the study using QCloud (Chiva et al. 2018).

For data analysis, extracted ion chromatograms corresponding to all monitored transitions were manually inspected using Xcalibur Qual Browser (Thermo Fisher Scientific). For shotgun proteomic datasets, peptide identification was performed using Proteome Discoverer (version 3.1) with the reviewed human proteome database from UniProt (Proteome ID: UP000005640; downloaded in October 2025), supplemented with custom variant-containing protein sequences and a common contaminant database. Database searches were carried out using the MASCOT search engine (server version 2.6.2.0), applying a precursor mass tolerance of 10 ppm and fragment ion tolerances of 0.02 Da or 0.6 Da, depending on the acquisition settings. Proteolytic specificity was defined according to the enzyme used (trypsin, GluC, or LysC), allowing up to two missed cleavages. Carbamidomethylation of cysteine residues was set as a fixed modification, while methionine oxidation and protein N-terminal acetylation were included as variable modifications. Peptide and protein identifications were filtered using a false discovery rate (FDR) of <1% at the peptide-spectrum match (PSM), peptide, and protein levels. Known contaminants were removed from downstream analyses. Protein abundance values were log₂-transformed, and normalization was performed based on the mean sample abundance to reduce technical variability associated with differences in sample loading (Aguilan et al. 2020). For PRM experiments, peptide peak areas were quantified using the open-source software Skyline (daily version 26.1) (MacLean et al. 2010). Transitions were selected based on fragment ion intensity, interference assessment, and reproducibility across replicates. Only transitions consistently detected across replicates were retained, while inconsistent or interfered transitions were manually excluded. Four fragment ions per peptide were ultimately used for quantification. PRM data were normalized using total ion current (TIC), and the ratios between alternative (Alt) and reference (Ref) peptide peak areas were calculated and log-transformed prior to statistical analysis. The complete shotgun proteomics and PRM datasets were deposited in the ProteomeXchange Consortium through the PRIDE partner repository (Y et al. 2025).

## Data access

The RNA-sequencing data developed for this study have been deposited. The BioProject and associated SRA metadata are available at: https://dataview.ncbi.nlm.nih.gov/object/PRJNA1456805?reviewer=1tcdr9orjfa28bp5vrggoo3 cp8 in read-only format.

The shotgun proteomics data were deposited in the PRIDE repository. The ProteomeXchang ID reserved for the data is: **PXD077902**. The DOI for the data is: https://doi.org/10.6069/srka-cq38.

PRM data were deposited in PANORAMA (https://panoramaweb.org/RPE.url). The reviewer account details are the following: Password: t7=AsLvFzyaVQg

## Competing interest statement

The authors declare no competing interests.

## Acknowledgements

This work was supported by Fondazione AIRC under the grant IG #25849 “Mining common genetic variants impacting on allele-specific translation and cancer risk” to A.I. A.L. and M.H.H. were supported by Short-term scientific mission fellowships by the “Translation Control in Cancer” European Network”, Translacore Cost Initiative (CA21154). This work was also partially supported by the initiative “*Dipartimenti di Eccellenza* 2023-2027 (Legge 232/2016)” funded by the MUR, and by FESR 2023 – “Sostegno alle Infrastrutture di Ricerca”. The European Regional Development Fund (ERDF) 2014-2020 POR P.A. Trento supports the CIBIO Mass Spectrometry and High-Throughput Screening Core Facilities at the University of Trento.

## Authors’ contributions

A.R., and A.I. conceptualized and supervised this work. L.A., M.H.H., D.P., E.D., and R.B. performed experiments and obtained the data used in this study. I.M. F.M., and R.B. performed data analysis. L.A. A.I. wrote the original draft of this manuscript. L.A. D.P. R.B., E.D., A.R., and A.I. revised this manuscript. All authors approved the submission of this manuscript.

## Supplementary Figures

**Figure S1.**
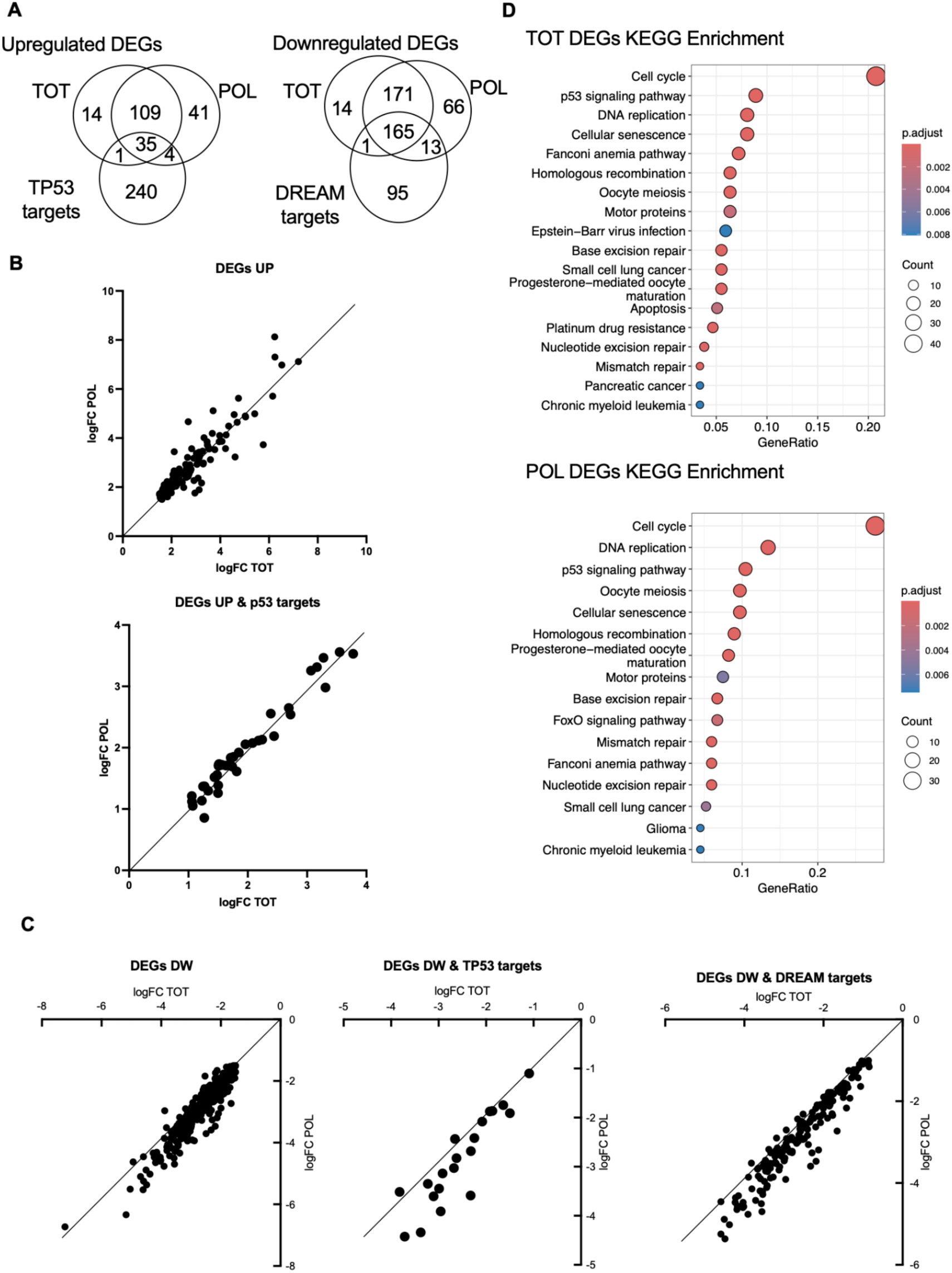
Nutlin treatment induces p53 and cell cycle arrest pathway responses in RPE-1 cells. **A)** Venn diagrams presenting the number of differentially expressed genes (DEGs) in response to Nutlin treatment in RPE1 cells and comparing the results obtained with total and polysome-associated mRNAs. Left diagram: upregulated DEGs. Genes were filtered for Log2 fold change above 1.5 and FDR<0.05. The majority of DEGs were common to both RNA fractions. The overlap with an curated, extensive list of TP53 target genes derived from the Fisher’s lab, is shown. Right diagram: downregulated DEGs. Genes were filtered for Log2 fold change below −1.5 and FDR<0.05. The majority of DEGs were common to both RNA fractions and there was a substantial overlap with a curated list of DREAM target genes, curated by the Engeland lab, that are indirectly repressed by p53. **B)** Comparison of the fold changes in total (X-axis) and polysomal (Y-axis) RNA fractions for upregulated DEGs. The majority of nutlin-induced gene expression changes are comparable in the transcriptome and the translatome, particularly for direct TP53 target genes (bottom panel). **C**) Comparison of the fold changes in total (X-axis) and polysomal (Y-axis) RNA fractions for downregulated DEGs. A stronger relative repression in polysomal RNA is apparent for the entire DEG list as well as for established TP53 (center panel) or DREAM targets (right), possibly consistent with a general inhibition of translation caused by Nutlin treatment (Attardi lab). **D**) KEGG semantic enrichment for the differentially expressed genes confirming the broad similarity among transcriptome and translatome data and the strong enrichment for cell cycle arrest and p53 activation signatures.

**Figure S2.**
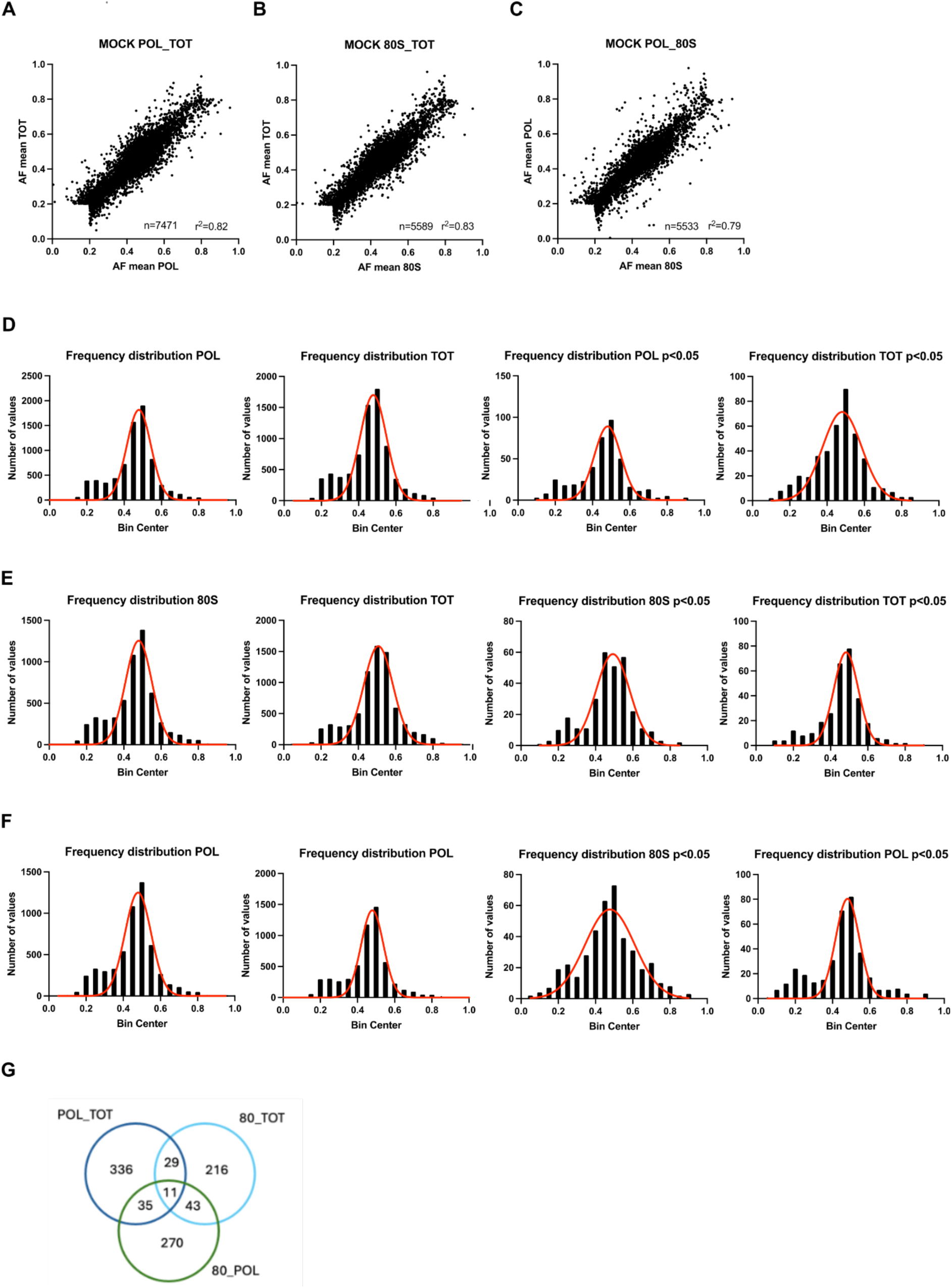
Allelic fraction distribution across 80S, POL and TOT comparisons. **A)** Distribution of allelic fraction (AF) values across the 7471 analyzable SNPs when comparing polysome and total-bound mRNAs displayed as a scatter plot for correlative analyses. **B)** Distribution of allelic fraction (AF) values across the 5589 analyzable SNPs when comparing 80S and total-bound mRNAs displayed as a scatter plot for correlative analyses. **C)** Distribution of allelic fraction (AF) values across the 5533 analyzable SNPs when comparing 80S and polysome-bound mRNAs displayed as a scatter plot for correlative analyses. **D)** Distribution of allelic fraction values across analyzable SNPs and tranSNPs (p < 0.05) for polysome and total-bound mRNAs. **E)** Distribution of allelic fraction values across analyzable SNPs and tranSNPs (p < 0.05) for 80S and total-bound mRNAs. **F)** Distribution of allelic fraction values across analyzable SNPs and tranSNPs (p < 0.05) for 80S and polysome-bound mRNAs. **G**) Overlap between TranSNP lists filtered on nominal p values (P<0.05).

**Figure S3.**
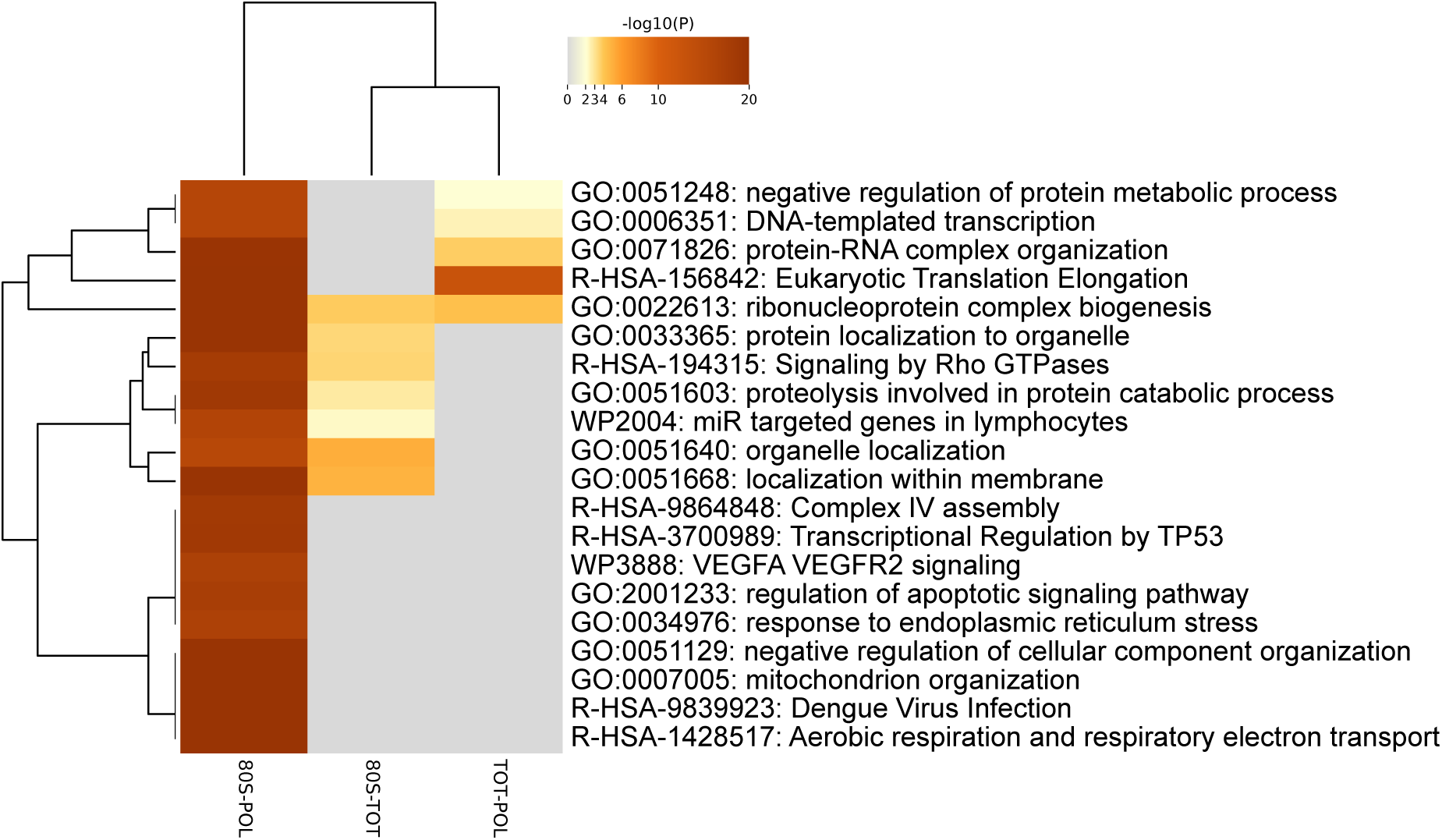
Semantic enrichment among differentially expressed genes across RNA fractions. Metascape heatmap view of the Gene Ontology Term enrichment for genes that were differentially expressed in the comparison between monosomal, polysomal, and total RNA fractions. See Methods for details.

**Figure S4.**
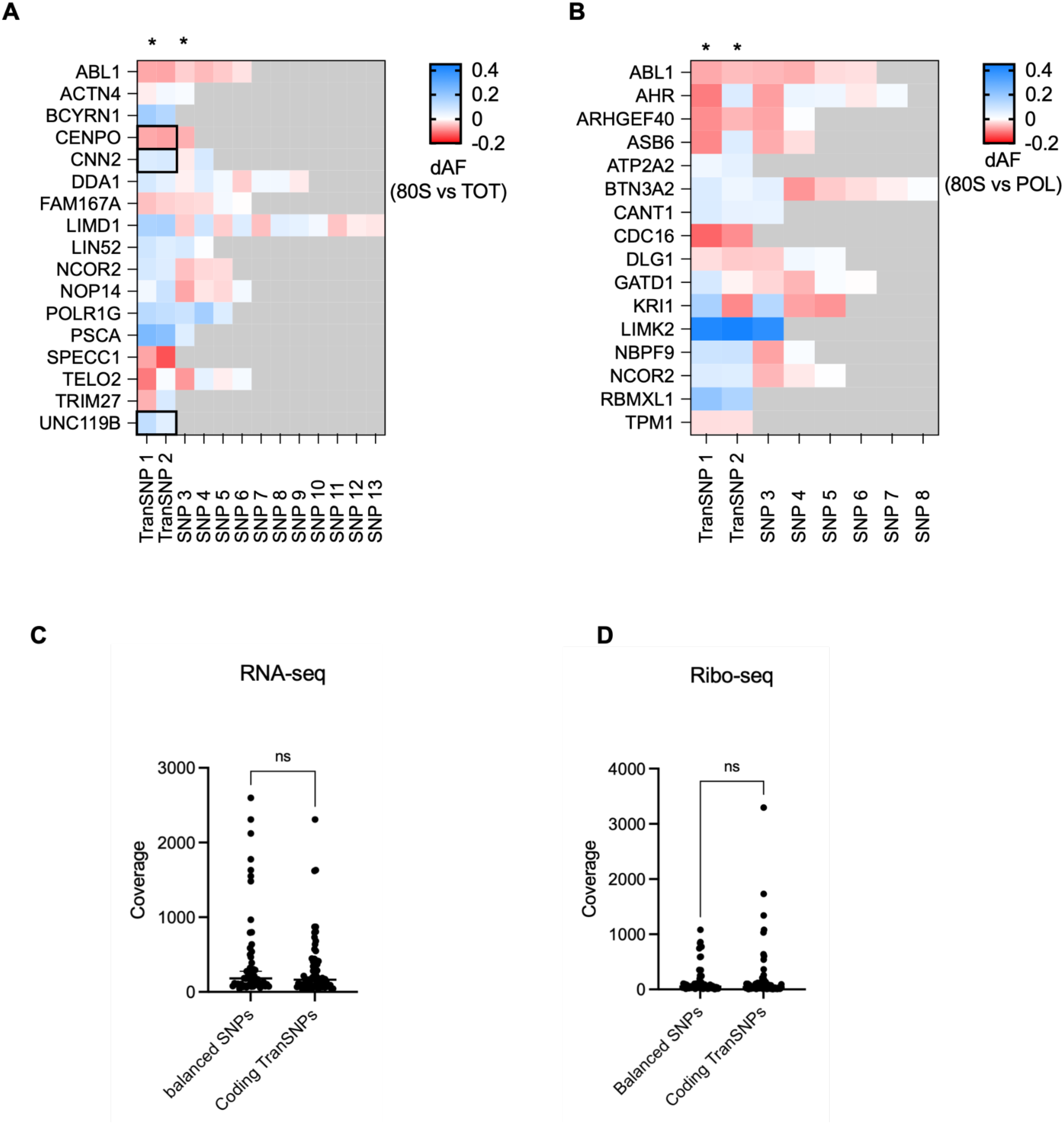
Extended delta allelic fraction data for 80S-associated haplotypes -related to Figure 2-. RNA-seq and Ribo-seq coverage -related to Figure 4. **A)** Delta allelic fraction data for the comparison between 80S and total RNA. The first two SNPs from the left in the heatmap that are also marked by an asterisk are the two identified tranSNPs (p<0.05). The following data is for additional SNPs that were analyzable in our dataset. The three black boxes mark cases where phasing data could be retrieved from the RNA-sequencing results and are in cis. **B**) Same as for panel A, but relative to the 80S versus polysome comparison. The data for the two tranSNPs for each haplotype are presented in Figure 2, along with the annotation of the phasing data. It is re-plotted here for easy comparison with the additional analyzable SNPs. For both panels, no additional phasing data could be retrieved from the RNA-sequencing. **C**, **D**) RNA-seq and Ribo-seq coverage for a control group of heterozygous RPE-1 SNPs not showing allelic imbalance, and the tranSNPs grouped into silent and missense alleles.

## Notes

### Competing Interest Statement

The authors have declared no competing interest.

## References

Aguilan JT, Kulej K, Sidoli S. 2020. Guide for protein fold change and: P-value calculation for non-experts in proteomics. Mol Omi 16: 573–582.

Andrysik Z, Galbraith MD, Guarnieri AL, Zaccara S, Sullivan KD, Pandey A, MacBeth M, Inga A, Espinosa JM. 2017. Identification of a core TP53 transcriptional program with highly distributed tumor suppressive activity. Genome Res 27: 1645–1657.

Biffo S, Ruggero D, Santoro MM. 2024. The crosstalk between metabolism and translation. Cell Metab 36.

Blandy A, Hopes T, Vasconcelos EJR, Turner A, Fatkhullin B, Agapiou M, Fontana J, Aspden JL. 2025. Translational activity of 80S monosomes varies dramatically across different tissues. Nucleic Acids Res 53.

Bolger AM, Lohse M, Usadel B. 2014. Trimmomatic: A flexible trimmer for Illumina sequence data. Bioinformatics 30: 2114–2120. http://www.ncbi.nlm.nih.gov/pubmed/24695404.

Chiva C, Olivella R, Borràs E, Espadas G, Pastor O, Solé A, Sabidó E. 2018. QCloud: A cloud-based quality control system for mass spectrometry-based proteomics laboratories. PLoS One 13.

Clamer M, Tebaldi T, Lauria F, Bernabò P, Gómez-Biagi RF, Marchioretto M, Kandala DT, Minati L, Perenthaler E, Gubert D, et al. 2018. Active Ribosome Profiling with RiboLace. Cell Rep 25: 1097–1108.e5.

Cook KB, Kazan H, Zuberi K, Morris Q, Hughes TR. 2011. RBPDB: A database of RNA-binding specificities. Nucleic Acids Res 39: D301–8.

Dobin A, Davis CA, Schlesinger F, Drenkow J, Zaleski C, Jha S, Batut P, Chaisson M, Gingeras TR. 2013. STAR: Ultrafast universal RNA-seq aligner. Bioinformatics 29: 15–21.

Fabbri L, Chakraborty A, Robert C, Vagner S. 2021. The plasticity of mRNA translation during cancer progression and therapy resistance. Nat Rev Cancer 21.

Fernández-Calero T, Davyt M, Perelmuter K, Chalar C, Bampi G, Persson H, Tosar JP, Hafstað V, Naya H, Rovira C, et al. 2020. Fine-tuning the metabolic rewiring and adaptation of translational machinery during an epithelial-mesenchymal transition in breast cancer cells. Cancer Metab 8.

Floor SN, Doudna JA. 2016. Tunable protein synthesis by transcript isoforms in human cells. Elife 5.

Hamadou M, Alunno L, Peroni D, Pancher M, Mazza F, Venturelli T, Belli R, Dassi E, Romanel A, Inga A. 2026. Polysomal Profiling Coupled to Allele-Specific Proteomics Reveals an EIF4H TranSNP Allele Possessing Higher mRNA Translation Potential. Mol Cell Proteomics 25.

Hamadou MH, Alunno L, Venturelli T, Valentini S, Dalfovo D, Lorenzini F, Mattivi A, Vigorito V, Grupelli GP, Matte A, et al. 2025. A TranSNP in the DDIT4 mRNA can impact its translation efficiency and modulate p53-dependent responses in cancer cells. BioRxiv https://ww.

Hermeking H. 2012. MicroRNAs in the p53 network: Micromanagement of tumour suppression. Nat Rev Cancer 12.

Heyer EE, Moore MJ. 2016. Redefining the Translational Status of 80S Monosomes. Cell 164: 757–769.

Huang E, Fu T, Zhang L, Yan G, Yamamoto R, Terrazas S, Nguyen TL, Gonzalez-Figueroa C, Khanbabaei A, Bahn JH, et al. 2025. Genetic variants affecting RNA stability influence complex traits and disease risk. Nat Genet 57.

Hughes CS, Moggridge S, Müller T, Sorensen PH, Morin GB, Krijgsveld J. 2019. Single-pot, solid-phase-enhanced sample preparation for proteomics experiments. Nat Protoc 14: 68–85.

Jia JJ, Lahr RM, Solgaard MT, Moraes BJ, Pointet R, Yang AD, Celucci G, Graber TE, Hoang HD, Niklaus MR, et al. 2021. MTORC1 promotes TOP mRNA translation through site-specific phosphorylation of LARP1. Nucleic Acids Res 49: 3461–3489.

Jia L, Mao Y, Ji Q, Dersh D, Yewdell JW, Qian SB. 2020. Decoding mRNA translatability and stability from the 5′ UTR. Nat Struct Mol Biol 27.

Koletsou E, Huppertz I. 2025. RNA-binding proteins as versatile metabolic regulators. NPJ Metab Heal Dis 3.

Koritzinsky M, Magagnin MG, Van Den Beucken T, Seigneuric R, Savelkouls K, Dostie J, Pyronnet S, Kaufman RJ, Weppler SA, Voncken JW, et al. 2006. Gene expression during acute and prolonged hypoxia is regulated by distinct mechanisms of translational control. EMBO J 25.

Kriachkov V, Ormsby AR, Kusnadi EP, McWilliam HEG, Mintern JD, Amarasinghe SL, Ritchie ME, Furic L, Hatters DM. 2023. Arginine-rich C9ORF72 ALS proteins stall ribosomes in a manner distinct from a canonical ribosome-associated quality control substrate. J Biol Chem 299.

Lee P, Chandel NS, Simon MC. 2020. Cellular adaptation to hypoxia through hypoxia inducible factors and beyond. Nat Rev Mol Cell Biol 21.

Li Z, Chen L. 2023. Predicting functional consequences of SNPs on mRNA translation via machine learning. Nucleic Acids Res 51.

Liang S, Bellato HM, Lorent J, Lupinacci FCS, Oertlin C, van Hoef V, Andrade VP, Roffé M, Masvidal L, Hajj GNM, et al. 2018. Polysome-profiling in small tissue samples. Nucleic Acids Res 46: E3.

Liao Y, Smyth GK, Shi W. 2019. The R package Rsubread is easier, faster, cheaper and better for alignment and quantification of RNA sequencing reads. Nucleic Acids Res 47.

Lin YJ, Huang LH, Huang CT. 2013. Enhancement of Heterologous Gene Expression in Flammulina velutipes Using Polycistronic Vectors Containing a Viral 2A Cleavage Sequence. PLoS One 8.

Lindström MS, Bartek J, Maya-Mendoza A. 2022. p53 at the crossroad of DNA replication and ribosome biogenesis stress pathways. Cell Death Differ 29.

Liu J, Lichtenberg T, Hoadley KA, Poisson LM, Lazar AJ, Cherniack AD, Kovatich AJ, Benz CC, Levine DA, Lee A V., et al. 2018. An Integrated TCGA Pan-Cancer Clinical Data Resource to Drive High-Quality Survival Outcome Analytics. Cell 173: 400–416.e11.

Love MI, Huber W, Anders S. 2014. Moderated estimation of fold change and dispersion for RNA-seq data with DESeq2. Genome Biol 15: 550. http://www.ncbi.nlm.nih.gov/pubmed/25516281.

Luo H, Kharas MG, Jaffrey SR. 2025. N6-Methyladenosine: an RNA modification as a central regulator of cancer. Nat Rev Cancer.

MacLean B, Tomazela DM, Shulman N, Chambers M, Finney GL, Frewen B, Kern R, Tabb DL, Liebler DC, MacCoss MJ. 2010. Skyline: An open source document editor for creating and analyzing targeted proteomics experiments. Bioinformatics 26.

Marcel V, Ghayad SE, Belin S, Therizols G, Morel AP, Solano-Gonzàlez E, Vendrell JA, Hacot S, Mertani HC, Albaret MA, et al. 2013. P53 Acts as a Safeguard of Translational Control by Regulating Fibrillarin and rRNA Methylation in Cancer. Cancer Cell 24: 318–330. http://www.ncbi.nlm.nih.gov/pubmed/24029231.

Marcel V, Van Long FN, Diaz JJ. 2018. 40 Years of Research Put P53 in Translation. Cancers (Basel) 10.

Mata J, Marguerat S, Bähler J. 2005. Post-transcriptional control of gene expression: A genome-wide perspective. Trends Biochem Sci 30.

Mitschka S, Mayr C. 2022. Context-specific regulation and function of mRNA alternative polyadenylation. Nat Rev Mol Cell Biol 23.

Mossmann D, Park S, Hall MN. 2018. mTOR signalling and cellular metabolism are mutual determinants in cancer. Nat Rev Cancer 18.

Orellana EA, Siegal E, Gregory RI. 2022. tRNA dysregulation and disease. Nat Rev Genet 23.

Panda A, Martindale J, Gorospe M. 2017. Polysome Fractionation to Analyze mRNA Distribution Profiles. Bio-Protocol 7: e2126.

Rauniyar N. 2015. Parallel Reaction Monitoring: A Targeted Experiment Performed Using High Resolution and High Mass Accuracy Mass Spectrometry. Int J Mol Sci 16: 28566–28581.

Rizzotto D, Zaccara S, Rossi A, Galbraith MD, Andrysik Z, Pandey A, Sullivan KD, Quattrone A, Espinosa JM, Dassi E, et al. 2020. Nutlin-Induced Apoptosis Is Specified by a Translation Program Regulated by PCBP2 and DHX30. Cell Rep 30: 4355–4369.e6.

Robert F, Pelletier J. 2018. Exploring the Impact of Single-Nucleotide Polymorphisms on Translation. Front Genet 9.

Robichaud N, Sonenberg N, Ruggero D, Schneider RJ. 2019. Translational control in cancer. Cold Spring Harb Perspect Biol 11.

Ruggero D. 2013. Translational control in cancer etiology. Cold Spring Harb Perspect Biol 5. http://www.ncbi.nlm.nih.gov/pubmed/22767671.

Schneider C, Erhard F, Binotti B, Buchberger A, Vogel J, Fischer U. 2022. An unusual mode of baseline translation adjusts cellular protein synthesis capacity to metabolic needs. Cell Rep 41.

Schug J. 2008. Using TESS to predict transcription factor binding sites in DNA sequence. Curr Protoc Bioinforma unit 2.6.

Tanenbaum ME, Stern-Ginossar N, Weissman JS, Vale RD. 2015. Regulation of mRNA translation during mitosis. Elife 4.

Thoreen CC, Chantranupong L, Keys HR, Wang T, Gray NS, Sabatini DM. 2012. A unifying model for mTORC1-mediated regulation of mRNA translation. Nature 485: 109–113.

Tian B, Yu S, Zhang Q. 2026. Regulation of gene expression by alternative polyadenylation in health and disease. Nat Rev Genet.

Tiu GC, Kerr CH, Forester CM, Krishnarao PS, Rosenblatt HD, Raj N, Lantz TC, Zhulyn O, Bowen ME, Shokat L, et al. 2021. A p53-dependent translational program directs tissue-selective phenotypes in a model of ribosomopathies. Dev Cell 56: 2089–2102.e11.

Tsai A, Kornberg G, Johansson M, Chen J, Puglisi JD. 2014. The dynamics of SecM-induced translational stalling. Cell Rep 7.

Valentini S, Gandolfi F, Carolo M, Dalfovo D, Pozza L, Romanel A. 2022. Polympact: Exploring functional relations among common human genetic variants. Nucleic Acids Res 50: 1335–1350.

Valentini S, Marchioretti C, Bisio A, Rossi A, Zaccara S, Romanel A, Inga A. 2021. TranSNPs: A class of functional SNPs affecting mRNA translation potential revealed by fraction-based allelic imbalance. iScience 24.

Volpe E, Colantoni A, Corda L, Di Tommaso E, Pelliccia F, Ottalevi R, Guarracino A, Licastro D, Faino L, Capulli M, et al. 2025. The reference genome of the human diploid cell line RPE-1. Nat Commun 16.

Wu Q, Bazzini AA. 2023. Translation and mRNA Stability Control. Annu Rev Biochem 92.

Xiang JS, Schafer DM, Rothamel KL, Yeo GW. 2024. Decoding protein–RNA interactions using CLIP-based methodologies. Nat Rev Genet 25.

Xiong Y, Wang A, Kang Y, Shen C, Hsieh CY, Hou T. 2025. mRNABERT: advancing mRNA sequence design with a universal language model and comprehensive dataset. Nat Commun 16.

Y P-R, C B, DJ K, S K, J B, S H, NS J, A P, M W, S W, et al. 2025. The PRIDE database at 20 years: 2025 update. Nucleic Acids Res 53: D543–D553.

Yoshikawa H, Larance M, Harney DJ, Sundaramoorthy R, Ly T, Owen-Hughes T, Lamond AI. 2018. Efficient analysis of mammalian polysomes in cells and tissues using ribo mega-SEC. Elife 7.

Zhao BS, Roundtree IA, He C. 2016. Post-transcriptional gene regulation by mRNA modifications. Nat Rev Mol Cell Biol 18.

Zheng D, Persyn L, Wang J, Liu Y, Ulloa-Montoya F, Cenik C, Agarwal V. 2025. Predicting the translation efficiency of messenger RNA in mammalian cells. Nat Biotechnol.

Zhou Y, Zhou B, Pache L, Chang M, Khodabakhshi AH, Tanaseichuk O, Benner C, Chanda SK. 2019. Metascape provides a biologist-oriented resource for the analysis of systems-level datasets. Nat Commun 10.

